# TENM4 is an essential transduction component for touch

**DOI:** 10.1101/2024.10.10.617546

**Authors:** Mohammed A. Khallaf, Angela Tzu-Lun Huang, Letizia Dalmasso, Sampurna Chakrabarti, Rowena Groeneveld, Yinth Andrea Bernal Sierra, Jonathan Alexis Garcia-Contreras, Emilie Nello, Anja Schütz, Valérie Bégay, Alessandro Santuz, Sarah Hedtrich, Wei Zhong, Christina Schiel, Steven J. Middleton, Oliver Popp, Niccolò Zampieri, Philipp Mertins, Severine Kunz, Gary R. Lewin

## Abstract

Gentle touch is conveyed to the brain by fast-conducting sensory fibers. Mechanosensitive ion channels at the terminals of these neurons are thought to be gated by extracellular tethers that transmit force from the surrounding matrix to the channel complex, but the molecular identity of such tethers has remained unknown. Here, we identify Teneurin-4 (TENM4) as an essential extracellular linker protein for mechanotransduction in mechanoreceptors. Sensory neuron–specific deletion of *Tenm4* in mice caused profound touch insensitivity, while acute and reversible proteolytic disassembly of TENM4 at sensory endings confirmed its direct role in force transduction. Ultrastructural analyses revealed TENM4 localization to filamentous structures at the neurite–laminin interface, defining it as a structural component of the mechanosensory tether. These findings identify TENM4 as a core element of fast somatic sensation and provide molecular insight into how extracellular forces are coupled to ion channel activation.

## INTRODUCTION

All organisms, from bacteria to humans, react within a fraction of a second to mechanical stimuli.^1,2^ In all branches of life, a molecularly diverse range of mechanosensitive ion channels are rapidly gated by force to initiate signaling.^2^ The fastest and most sensitive mechanosensitive channels detect nanometer scale movements that enable hearing and touch perception.^3–5^ Interestingly, mechanosensitive channels in fruit flies, worms and vertebrate hair cells require intracellular or extracellular tether proteins that relay force to the channel.^5–11^ Vertebrate touch sensation also requires mechanosensitive ion channels, of which PIEZO2 and ELKIN1 share roles in mechanosensation.^4,12,13^ The skin is supplied with a rich variety of molecularly and physiologically distinct mechanoreceptors,^14^ the cell bodies of which reside in the dorsal root ganglia (DRG). Despite their functional diversity, all mechanoreceptors transduce mechanical stimuli into electrical signals at specialized neuroglial endings in the skin.^15,16^ There are several lines of evidence that extracellular proteins at these neuroglial endings are directly involved in transferring force to mechanically gated channels. First, mechanically activated currents in sensory neurons can be acutely ablated by proteases, including blisterase, an enzyme that specifically cleaves furin consensus sites.^17,18^ Mechanosensitive currents are restored 24 hours after protease treatment in cultured sensory neurons, suggesting that the protease target is synthesized by sensory neurons.^17^ In the skin, protease treatment of the receptive fields of individual mechanoreceptors rapidly abolishes mechanosensitivity indicating an effect at the neuroglial ending.^17^ Protease treatment was also associated with the loss of tether-like structures synthesized by sensory neurons.^17^ These data, although suggestive, did not identify the molecular nature of the extracellular protein(s) that are necessary for transduction. Here, we identify TENM4 as structural component in mechanotransduction complexes the integrity of which is necessary for transduction in essentially all cutaneous mechanoreceptors.

## RESULTS

### Screening for tether candidates

Blisterase cleaves a furin consensus site RX(K/R)R, which was predicted to occur in at least 11,302 mouse proteins; but only a small minority of these possess putative cleavage sites located in protease accessible extracellular domains. Candidates with predicted extracellular domains were selected using the Database for Annotation, Visualization, and Integrated Discovery (DAVID).^19^ Using gene paint (www.genepaint.org),^20^ we selected 34 of these genes for further analysis based on the presence of in situ hybridization signals in the developing and adult DRG. Sympathetic neurons located in the superior cervical ganglion (SCG) do not possess mechanically activated currents (MA currents).^17,21^ We thus reasoned that our candidate(s) should be expressed in the DRG, but not in the SCG. Using qPCR, we could not detect the mRNA for six of the 34 genes in the SCG, but all showed robust mRNA expression in the DRG (Table S1). The six genes coded for the following proteins: Teneurin-4 (TENM4), FRAS1 Related Extracellular Matrix Protein 2 (FREM2), Proprotein Convertase Subtilisin/Kexin Type 5 (PCSK5), Scribble Homolog (SCRIB), Alpha-Tectorin (TECTA), and Slit Guidance Ligand 1 (SLIT1). We used antibodies directed against the six candidates and obtained signals in cultured DRG neurons for each candidate, but not in SCG cultures, validating the results of the qPCR experiment (Figure S1A). We chose to study TENM4 and FREM2 first, partly because these proteins had the largest extracellular domains of the candidates (mouse proteins 2771 aa and 3160 aa, respectively), consistent with the large tether structures we had previously observed.^17,22^ We generated two mouse strains with floxed *Frem2* and *Tenm4* alleles (Figures S2A and S2B) as both *Frem2* and *Tenm4* appear to be essential genes during embryonic development.^23,24^

Sensory neuron-specific conditional knockouts were created using an adeno-associated virus that is neurotropic for DRG neurons carrying a Cre/GFP cassette (AAV-PHP.S-Cre-eGFP).^25,26^ We examined *Frem2^fl/fl^* mice 4-6 weeks after virus application, a time point chosen to allow sufficient Cre-mediated excision.^26^ Paw withdrawal thresholds to von Frey hair stimulation, responses to cotton swab and brush stimuli indicated no loss of touch sensitivity in these mice (Figure S1B). We thus turned our attention to the potential role of TENM4 in sensory mechanotransduction.

### Mechanoreceptors express the TENM4 protein

TENM4 is a member of the Teneurin family, consisting of four proteins in vertebrates, which are evolutionarily conserved type II membrane proteins known to be essential during neural development.^27,28^ Teneurins are dimeric proteins and there is some evidence that hetero-dimers between two different Teneurin proteins may be formed.^29^ We identified a commercially available TENM4 antibody directed against the intracellular N-terminus, which is divergent between the four mouse Teneurins (Anti-TENM4-N).^30^ But we also generated an antibody against a large recombinant extracellular fragment, which included the extracellular compact YD shell region of the protein (Anti-TENM4-C). We first looked for TENM4 at the neuroglial endings of a variety of mechanoreceptor types in the skin where touch stimuli are transduced (Figure 1A),^15,16^ and found strong immunolabeling of sensory endings that were also positive for neurofilament-200 (NF200^+^), a marker of all myelinated sensory fibers (Figure 1B). Sensory end-organs innervated by rapidly-adapting mechanoreceptors (RAMs) like Meissner corpuscles and hair follicles were all strongly positive for TENM4 (Figure 1B). Antibodies against S100β were used to label terminal Schwann cells that make up the Meissner corpuscle and are closely associated with lanceolate endings around hair follicles,^15,31^ however TENM4 labelling was not seen in these cells (Figure 1B). We found that slowly-adapting mechanoreceptors (SAMs) positive for NF200^+^ that innervate CK20^+^ Merkel cells were also co-labelled with TENM4 (Figure 1B). Using the same strategy as for *Frem2*, we generated mice with a sensory neuron specific deletion of *Tenm4* (Figures 2A and S2B) and saw essentially no TENM4 labeled sensory endings associated with RAM or SAM innervated end-organs (Figure 1B), demonstrating the specificity of the antibody (Anti-TENM4-N). We also detected TENM4^+^ axons in wild type (WT) skin that terminated as free nerve endings (Figure S1C). Thus, TENM4 appeared to be a good marker of myelinated sensory endings including low threshold mechanoreceptors (LTMRs).

**Figure 1.**
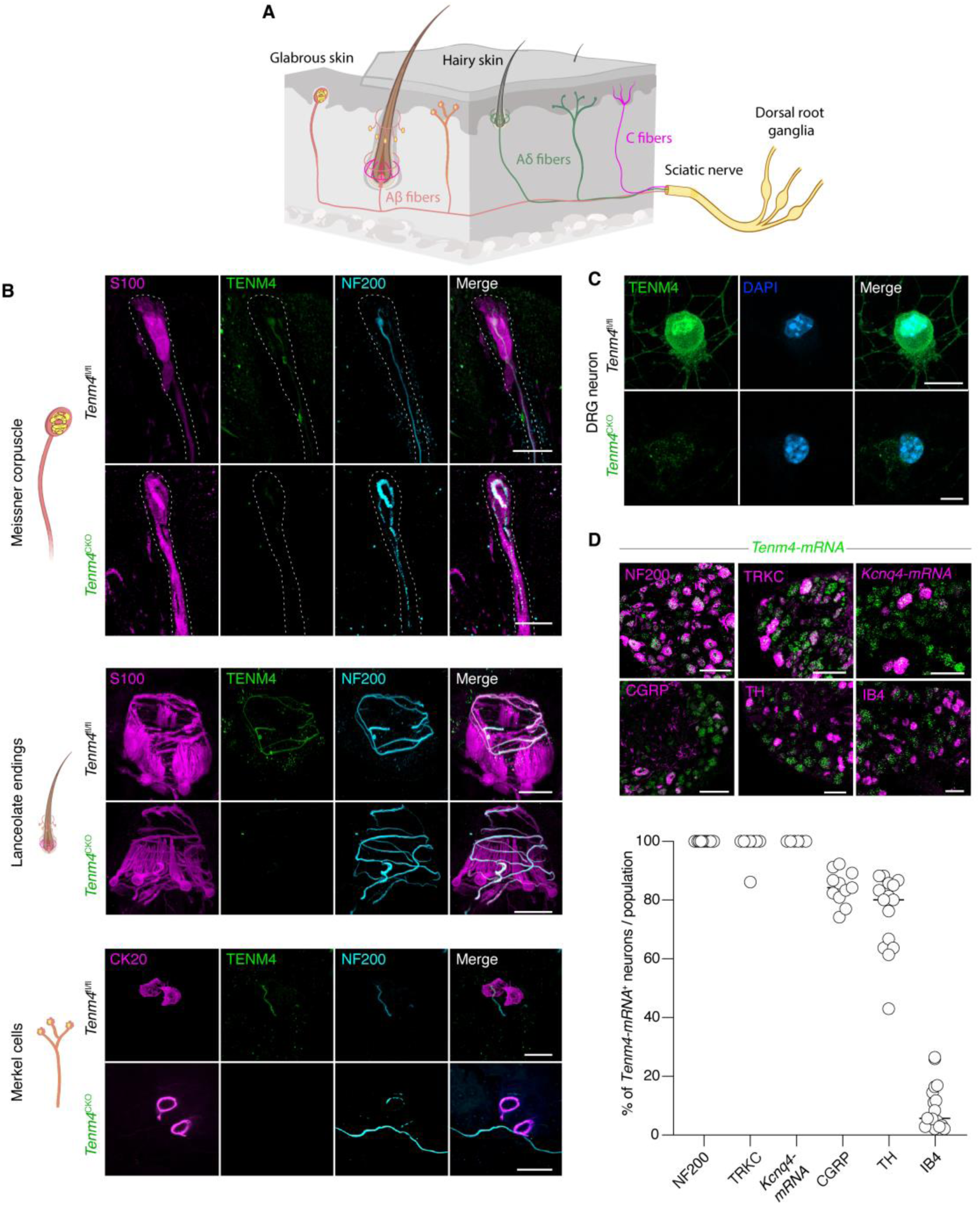
TENM4 is present at sites of transduction in mechanoreceptors. (**A**) Schematic of mechanoreceptor and nociceptor endings in hairy and glabrous skin. (**B**) TENM4+ and NF200+ sensory endings associated with rapidly-adapting mechanoreceptors (RAMs) like Meissner’s corpuscles, lanceolate endings around hair follicles and slowly adapting mechanoreceptors (SAMs) innervating CK20+ Merkel cells. S100+ terminal Schwann cells were not TENM4+. TENM4 immunostaining was largely absent from RAM and SAM endings in *Tenm4^CKO^* mice. Scale bars 20μm. (**C**) TENM4+ staining was absent in cultured sensory neurons from *Tenm4^CKO^* mice. Scale bar 10μm. (**D**) Representative DRG sections showing double labelling of *Tenm4* smFISH probe with NF200, TRKC, *Kcnq4-mRNA*, CGRP, TH or IB4, Scale bar 100μm. Bottom shows quantification Tenm4+ sensory neurons co-labeled with different sensory markers. For each co-labelling 2 male and 3 female mice were used and more than 700 neurons counted.

**Figure 2.**
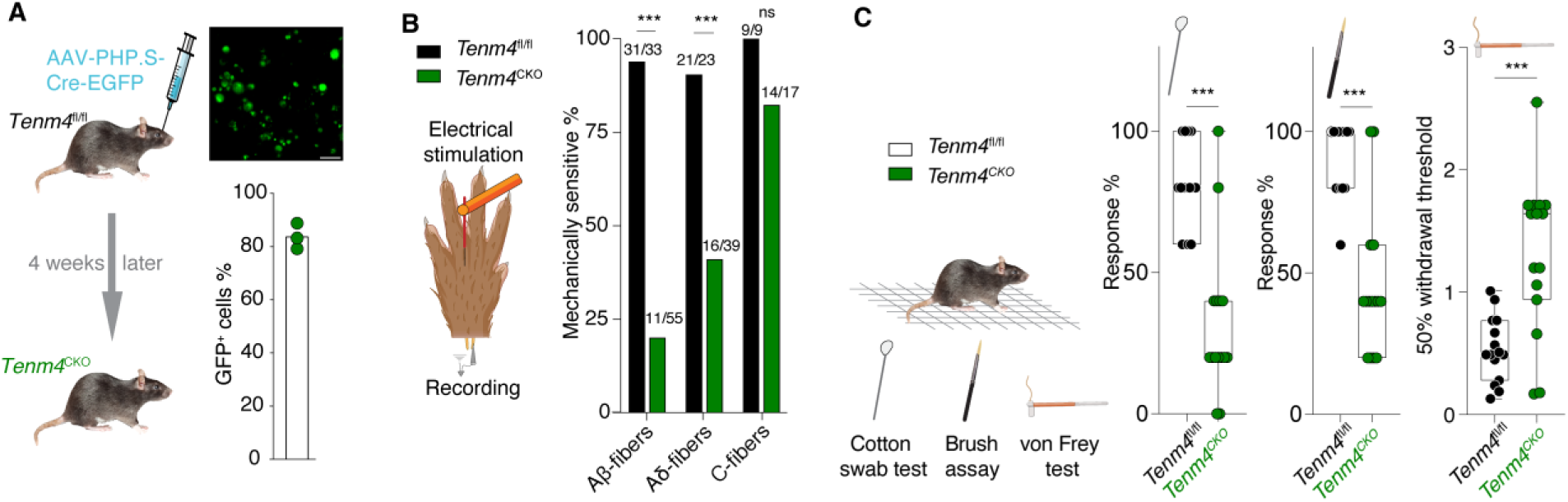
TENM4 is required for mechanoreceptor function and touch sensation. (**A**) Left, schematic for generation of *Tenm4^CKO^* mice via retro-orbital injection of AAV-PHP.S-Cre-EGFP of *Tenm4^fl/fl^* mice. Top right, representative image of DRG sensory neurons from *Tenm4^CKO^* mice 4 weeks after being transduced with AAV-PHP.S-Cre-EGFP. Scale bar is 50 µm. Bottom right, quantification of the percentage of GFP positive cultures DRG neurons after viral transduction. Dots represendata from 3 mice. (**B**) Electrical search technique (schematic) used to identify saphenous fibers with and without a mechanosensitive receptive field. *Tenm4^CKO^* mice exhibited a substantial increase in Aβ-fibers and Aδ-fibers without a mechanosensitive receptive field, C-fibers remained unaffected. Significance calculated with Fisher’s exact test, data from 5 *Tenm4^fl/fl^* (control) and 10 *Tenm4^CKO^* mice. (**C**) Behavioral tests on touch evoked behaviors. *Tenm4^CKO^* mice exhibited reduced responses to a cotton swab and a dynamic brush. Additionally, *Tenm4^CKO^* mice exhibited three-fold higher withdrawal thresholds and significant deficits in mean force required for a 50% withdrawal using von Frey hairs, compared to controls. Data were collected from 15 *Tenm4^fl/fl^*and 15 *Tenm4^CKO^* mice. Significance calculated with Student’s T-Test.

Our validated antibody revealed the presence of TENM4 in large diameter DRG neurons in culture as well as a specific localization to the mechanoreceptor endings associated with specific end-organs in the skin (Figures 1B,C and S1E). We next analyzed *Tenm4* expression by single-molecule fluorescence in situ hybridization (smFISH) and assessed its co-localization with markers of nociceptors and mechanoreceptors in DRG sections (Figure 1D). Consistent with our results from the skin, all NF200^+^ cells were also *Tenm4^+^*. However, many nociceptive sensory neurons positive for calcitonin-gene related peptide (CGRP) were also co-labeled with *Tenm4* (Figure 1D). TRKC and *Kcnq4* expression are highly specific to RAMs,^32,33^ and all these cells were also positive for *Tenm4* (Figure 1D). Small sensory neurons positive for tyrosine hydroxylase (TH), a marker of low threshold tactile C-fibers^34^, were also largely positive for *Tenm4*. However, we saw very little, or no overlap with nociceptor markers such as isolectin-B4 (IB4) (Figure 1D). As expected, the TENM4 signal was largely absent in cultured sensory neurons and sections from *Tenm4^CKO^*mice (Figures 1C and S1E). Together, these data indicate that TENM4 is selectively present in virtually all mechanoreceptors including nociceptors with myelinated axons, suggesting its involvement in touch and rapid pain sensation.

### *Tenm4* deletion abolishes mechanosensitivity and causes sensory deficits

We next tested whether TENM4 is necessary for the mechanosensory function of identified mouse mechanoreceptors using *Tenm4^CKO^* mice (Figure 2A). Four weeks after retro-orbital injection of AAV-PHP.S-Cre-EGFP virus in *Tenm4^fl/fl^*mice we cultured sensory neurons from the DRG and found that ∼80% showed green fluorescence. Thus, the vast majority of the sensory neurons had probably undergone Cre-mediated excision at the *Tenm4* locus (*Tenm4^CKO^*mice). From the same mice we used an ex vivo saphenous skin-nerve preparation to trace the trajectory of single units by means of an electrical stimulus until the point of exit from the nerve branch. Normally, single identified Aβ- and Aδ-fibers always have a mechanosensitive receptive field in the skin near to the nerve exit point, which we confirmed here in control mice (*Tenm4^fl/fl^*) (Figure 2B).^4,12,35,36^ However, blinded recordings made from *Tenm4^CKO^* mice revealed that >75% of the Aβ-fibers and 60% of Aδ-fibers had no detectable mechanosensitive receptive field (Figure 2B). Recordings from C-fiber nociceptors revealed no significant differences in the presence of mechanosensitive receptive fields between control and *Tenm4^CKO^*mice (Figure 2B). The deletion of the *Tenm4* gene in the majority of DRG neurons did not lead to axon loss as determined with transmission electron microscopy (Figure S3A) nor to any changes in axonal conduction velocity (Figures S3B and S3C).

Expression of the virally expressed GFP marker in sensory neurons of *Tenm4^CKO^* mice indicated that around 80% of the cells were transduced 4 weeks post-injection (Figure 2A). However, the presence of Cre protein will not always lead to complete gene excision 4 weeks after transduction. We therefore asked whether the remaining mechanosensitive fibers may have impaired function due to incomplete ablation of the TENM4 protein in sensory neurons. We thus examined the stimulus response characteristics of the remaining non-silenced sensory fibers identified using a mechanical search stimulus in both glabrous and hairy skin.^16,37^ We used a 20Hz vibrating stimulus with continuously increasing amplitude to stimulate RAMs in hairy and glabrous skin. We found that all RAMs displayed significantly reduced firing rates in *Tenm4^CKO^* mice compared to control mice (Figure S3D). Velocity sensitivity was also measured using ramp stimuli of different velocities and spike rates to these stimuli were also significantly impaired in RAMs from *Tenm4^CKO^* mice (Figure S3E). Low threshold mechanoreceptors with Aδ-fiber conduction velocities, so called D-hair receptors, showed significantly higher vibration thresholds in *Tenm4^CKO^*mice compared to controls (Figure S3D). Merkel cells are innervated by slowly adapting mechanoreceptors (SAMs) and are found in hairy and glabrous skin. Stimulus response functions calculated with a series of 10s long ramp and hold stimuli showed that SAMs fired significantly less and had higher mechanical thresholds for activation in *Tenm4^CKO^* mice compared to controls (Figures S3E and S3F). We also recorded from Aδ mechanonociceptors (AMs) which detect sustained high threshold mechanical stimuli and contribute to fast mechanical pain sensation.^38^ In *Tenm4^CKO^* AMs displayed reduced sustained firing to high forces and stimulus response functions were reduced, but this was not significantly different compared to controls (Figures S3F and S3G). Taken together, our data show that normal levels of TENM4 are necessary for the mechanosensory function of almost all cutaneous sensory receptors, with the exception of unmyelinated C-fibers (Figure 2B). We predicted that the dramatic functional loss of sensory neuron mechanosensitivity in *Tenm4^CKO^*should lead to profound behavioral deficits, especially to touch. Indeed, *Tenm4^CKO^*mice displayed substantially reduced responses to a cotton swab and dynamic brush stimuli (Figure 2C). Additionally, paw withdrawal thresholds to a series of von Frey hairs were substantially elevated in *Tenm4^CKO^* mice (Figures 2C and S4A). Sensory deficits varied in degree between individuals probably due to differences efficiency of gene excision, but in many cases, animals completely failed to respond to mechanical stimuli indicating a profound loss of sensation (Figures 2C and S4A).

We confirmed these results using a complementary strategy in which we generated mice in which the *Tenm4^fl/fl^* alleles were crossed with mice carrying an *Advillin-CreERT2* allele^4^ allowing the inducible deletion of *Tenm4* only in sensory neurons following tamoxifen treatment. Around 4 weeks after tamoxifen injections *Avil^CreERT2^:Tenm4^fl/fl^*mice were tested for paw withdrawal using a series of von Frey hairs of increasing force. As for virus injected mice, these treated animals displayed profoundly elevated paw withdrawal thresholds (Figure S4C). Using a radiant heat stimulus, we measured thermal thresholds in both *Avil^CreERT2^:Tenm4^fl/fl^*and virus injected *Tenm4^CKO^* mice and found no significant deficits (Figures S4B and S4C). Loss of touch sensation may also lead to altered coordination and indeed we noted that *Tenm4^CKO^* mice showed hindlimb clasping that was rarely seen in control mice (Figure S4D). We found that essentially all Parvalbumin positive DRG soma were also stained positive for TENM4 (Figure S4E), Parvalbumin is a marker for proprioceptors, although it also probably found in some cutaneous sensory neurons.^39,40^ We generated conditional *Tenm4* mutant in which the gene was deleted in proprioceptors by crossing our *Tenm4^fl/fl^*mice with a *Parvalbumin-Cre* line (*Pv-Cre:Tenm4^fl/fl^* mice).^41^ We found that *Pv-Cre:Tenm4^fl/fl^*mice showed mild impairments in their ability to remain on a rotating wheel (rotarod) test compared to controls (*Tenm4^fl/fl^* mice) (Figure S4D). These deficits could be due to alterations in proprioceptors, but may also reflect the reduced input from cutaneous afferents that also contribute to normal proprioception.^39,40,42^ This prompted us to ask whether we could detect TENM4 protein at the endings of proprioceptors that innervate the muscle spindle. We visualized proprioceptor endings in muscle spindles from the mouse gastrocnemius muscle using *Pv^cre^;Rx3^Flpo^;Ai65D* mice^43^ (see methods). We failed to detect TENM4 signal in identified proprioceptor endings (Fig. S4F). It thus appears likely that motor coordination deficits seen following conditional *Tenm4* deletion were due to a loss in cutaneous sensory feedback in these animals. These data also suggest that not all sensory neurons that express TENM4 can transport the protein to the sensory end-organ.

### TENM4 cleavage abolishes touch sensitivity

Cre-mediated excision using AAV vectors leads to the removal of *Tenm4* over several weeks. TENM4 has well described roles in intercellular signaling,^30,44^ the alteration of which could have indirect effects on transduction. Therefore, we wished to directly test the idea that TENM4 forms part of a complex that facilitates force transfer from the extracellular matrix (ECM) to transduction channels. Ideally, specific cleavage of proteins linking the channel to ECM should abolish transduction.^17^ We thus developed a new technology that enabled us to specifically ablate TENM4 function within minutes. We named this method PreScission Protein Disassembly, and it is based on the fact that the plant virus derived PreScission protease cleaves a specific consensus sequence (LEVLFQ↓GP),^45^ that rarely occurs in the extracellular exposed residues of mouse proteins (Table S2). This method is conceptually similar to a method recently used in nematodes to genetically target proteolytic cleavage of transduction molecules.^10^ Using homologous recombination in mouse ES cells we inserted the PreScission cleavage sequence into *Tenm4*. We replaced 9 codons leading to the replacement of 7 amino acids which expose a PreScission sequence on the surface of the extracellular YD shell domain of TENM4.^44^ This manipulation was predicted to make TENM4 newly susceptible to cleavage by the PreScission enzyme (Figures 3A and S2C). Mice carrying the PreScission site in the *Tenm4* gene were bred, but out of 569 newborns from heterozygote crosses we obtained no homozygote animals. The failure to obtain homozygotes indicates that altering both *Tenm4* alleles likely causes embryonic lethality, consistent with previous studies.^30,46^ We thus decided to work with heterozygotes which we termed *Tenm4^KI/+^* mice. Considering the observed homozygote embryonic lethality, it was important to examine whether *Tenm4^KI/+^*mice show deficits in baseline sensory function. *Tenm4^KI/+^* mice exhibited uncleaved full-length TENM4 (Figure 3B), and showed no deficits in behavioral responses to mechanical and thermal stimuli (Figures S5A-C). We also made recordings from the saphenous nerve of *Tenm4^KI/+^* mice and, using the electrical search method, demonstrated that Aβ-fibers, Aδ-fibers and C-fibers exhibited normal mechanosensitivity, with unaltered stimulus response properties and normal axonal conduction velocities (Figures 3C and S6A). Thus, inclusion of mutant TENM4 protein into the mechanotransduction complex does not impact on baseline sensory function. Since TENM4 forms obligate homodimers,^29,44^ a stochastic mixing of mutated and wild type proteins would ensure that the vast majority of dimers would have at least one PreScission cleavable site. Indeed, we directly confirmed that the PreScission enzyme cleaves TENM4 in vivo. Western blots from glabrous skin revealed a 300 kDa band corresponding to uncleaved TENM4 in WT skin treated with PreScission protease. However, in skin samples taken from *Tenm4^KI/+^*mice treated with PreScission, we detected not only the full-length protein but also a ∼180 kDa fragment, consistent with the predicted size of the cleavage product (Figure 3D). Thus endogenous dimeric complexes at sensory endings consist of both WT and PreScission cleavable protein. We next applied the PreScission protease directly to the receptive fields of single identified mechanoreceptors to ask whether acute TENM4 ablation can rapidly abolish mechanosensitivity as we previously saw using the Blisterase enzyme.^17^ Once a mechanoreceptor was identified and classified,^37^ the receptive field was isolated with a metal ring and repeatedly stimulated every minute for up to 45 mins. We used a vibration stimulus (20Hz gradually increasing to an amplitude of 100 mN) for all LTMRs (RAMs, SAMs and D-hair receptors) or a 150mN amplitude ramp and hold stimulus for mechanonociceptors (AMs or C-fiber nociceptors). After baseline recordings for three minutes the receptive field was exposed to the PreScission enzyme (2 units/µL) in physiological ringer solution (Figure 3E). Protease treated LTMRs from WT mice showed remarkably robust spiking for the entire 45 mins with barely any diminution of mechanosensitivity (Figures 3E and 3F). In contrast, all LTMRs (RAMs, SAMs and D-hair receptors) recorded from *Tenm4^KI/+^*mice showed progressively decreasing spike responses to the constant mechanical stimuli starting just minutes after protease application, until no response was evoked between 30 and 45 minutes (Figure 3F). Complete loss of mechanosensitivity was also seen in RAMs innervating the Meissner’s corpuscles in glabrous skin (Figure 3F). For each mechanoreceptor examined we used an electrical stimulus at the end of the experiment and could show in every case that the electrically evoked spike was always still present, indicating a specific loss of mechanosensitivity. The rapid loss of LTMR mechanosensitivity could in theory be due to a toxic effect of the PreScission cleavage product on the mechanoreceptors. This idea was directly tested by adding the fluid containing the cleavage product from PreScission treated *Tenm4^KI/+^* skin to the isolated receptive fields of WT neurons (Figure S7A). The buffer containing the cleavage product had no effect on the mechanosensitivity of WT RAMs (Figure S7A).

**Figure 3.**
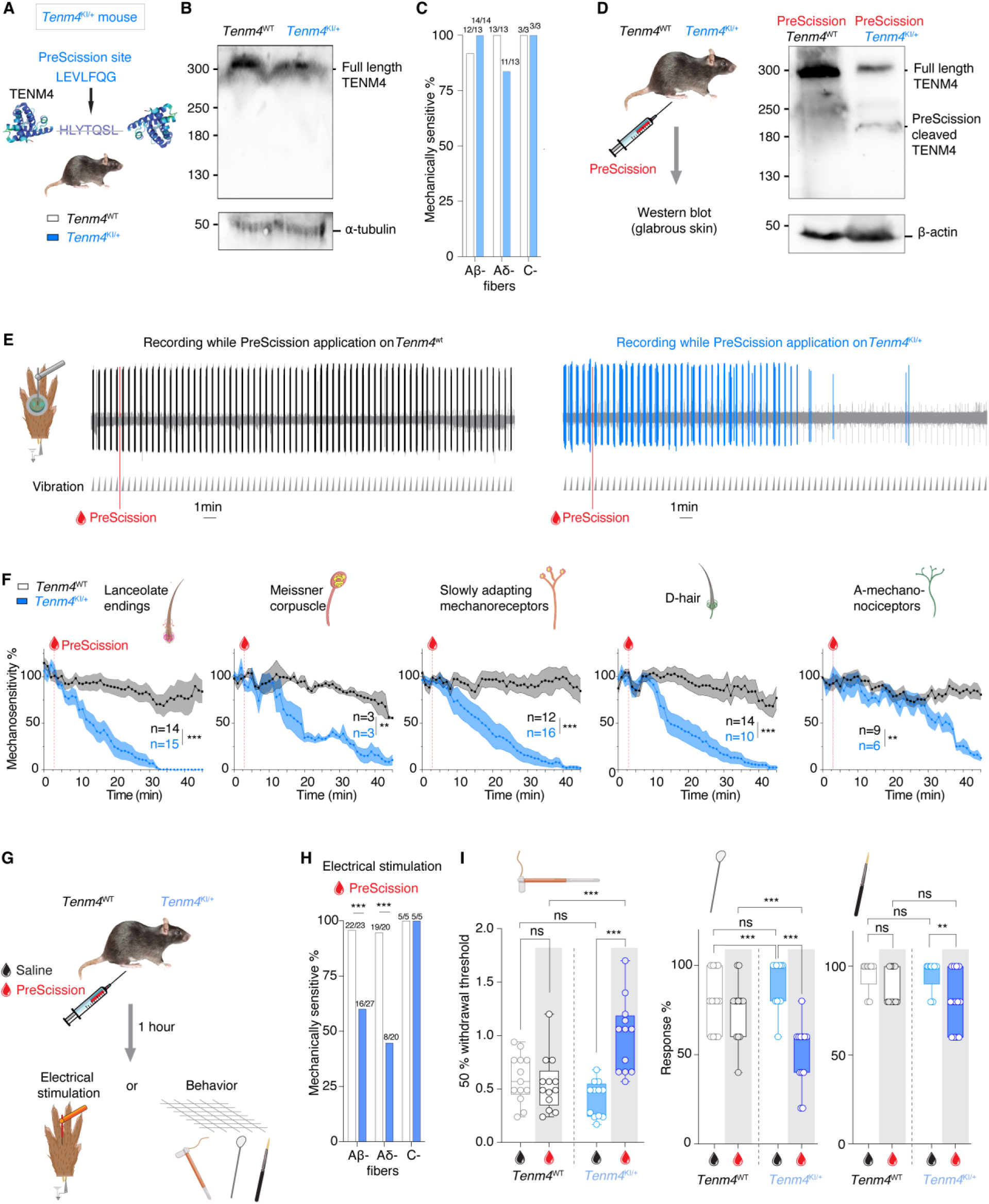
Rapid loss of mechanoreceptor function following TENM4 cleavage. (**A**) Generation of *Tenm4^KI/+^* mice: insertion of a 7 residue PreScission consensus sequence into genomic locus encoding the TENM4 extracellular domain. (**B**) western blot of endogenous TENM4 from glabrous skin revealed uncleaved full-length TENM4 in WT and *Tenm4^KI/+^* mice (**C**) Electrical search technique revealed that most Aβ, Aδ, and C-fibers retained mechanosensitivity in *Tenm4^KI/+^* mice as in WT mice. Significance calculated with Fisher’s exact test. Data collected from 4 WT (control) and 5 *Tenm4^KI/+^* mice. (**D**) Left, experimental design. Right, western blot of endogenous TENM4 from glabrous skin after PreScission protease treatment revealed uncleaved full-length TENM4 in WT mice, a cleaved product of the predicted size was detected in *Tenm4^KI/+^*mice. Loading control β-actin. (**E**) Representative example of a single unit (RAM) responding to repeated 20Hz vibration before and after application of PreScission to the receptive field in a WT (left) or *Tenm4^KI/+^*(right) mouse. Note the progressive decrease in stimulus induced firing following PreScission treatment only in *Tenm4^KI/+^* (right) mouse. (**F**) Mean mechanoreceptor responses plotted as a percentage of the starting value for all myelinated Aβ and Aδ types, before and after PreScission. Note mechanoreceptors in WT all retained mechanosensitivity (grey), but all mechanoreceptor types in *Tenm4^KI/+^* mice, progressively lost responsiveness during the 45 mins of observation (blue). (**G**) Experimental design. (**H**) Electrical search 1 hour after PreScission protease injections. Substantial and significant decrease in the proportion of Aβ and Aδ-fibers displaying mechanosensitivity in PreScission treated *Tenm4^KI/+^*, but not in WT mice. The proportion of mechanosensitive C-fibers remained unchanged in both groups. Significance calculated with Fisher’s exact test. Data collected from 4 WT and 4 *Tenm4^KI/+^* mice. (**I**) Behavioral assays used to assess the effects of TENM4 cleavage 1 hr after PreScission injection to the plantar hind paw. Mechanical withdrawal threshold measured with von Frey hairs were unchanged in WT mice treated with saline or PreScission. Additionally, *Tenm4^KI/+^* mice injected with PreScission showed decreased responses to both a cotton swab and a dynamic brush, no changes were observed in treated WT mice. Data collected from 13 WT and 12 *Tenm4^KI/+^* mice. Two-way repeated measures ANOVA followed by Sidak’s post-test. ns indicates p > 0.05; * p < 0.05; ** p < 0.01; ***; p < 0.001, means ± s.e.m.

Notably, we observed a slower, but robust loss of the ability of AMs in *Tenm4^KI/+^* to transduce force, reaching its maximum 45 mins after PreScission application (Figure 3F). However, in contrast to LTMRs, most AMs retained residual responses to intense mechanical stimuli at the end of the 45-minute observation period (5/6 tested). C-fiber nociceptors in WT and *Tenm4^KI/+^* mice retained functionality after exposure to PreScission protease (Figure S7B), indicating a fundamentally distinct transduction mechanism in AM nociceptors compared to C-fiber nociceptors.

We next performed behavioral and electrophysiological assays on mice subjected to subcutaneous injections of the PreScission enzyme into either the glabrous skin or hairy skin innervated by the saphenous nerve (Figure 3G). One hour after subcutaneous injections of the PreScission enzyme we observed a substantial and significant decrease in the proportion of Aβ- and Aδ-fibers with mechanosensitive receptive fields in *Tenm4^KI/+^* mice compared to WT mice in the saphenous nerve (Figure 3H). Not all mechanoreceptors lost their mechanosensitivity, likely because the subcutaneously injected enzyme did not reach the entire innervation territory of the saphenous nerve. However, no loss of C-fiber mechanosensitivity was observed in either genotype treated with the PreScission protease (Figures 3H and S7B). We next asked whether touch evoked behaviors which are routinely measured using stimuli to the glabrous skin are impacted by the application of the PreScission protease. In WT mice PreScission injections into the glabrous skin had no significant impact on paw withdrawal thresholds to von Frey stimuli, cotton swab or brush responses. However, *Tenm4^KI/+^* mice exhibited a substantial and significant increase in paw withdrawal thresholds as well as much reduced responses to cotton swab and brush stimuli (Figures 3I and S7C). Noxious heat withdrawal latencies were not significantly impacted after PreScission injection in *Tenm4^KI/+^* mice compared to saline injected controls (Figure S7D).

### TENM4 is essential for touch sensation

Cleavage of the TENM4 protein clearly leads to an acute loss of touch sensation in *Tenm4^KI/+^* mice, but in theory the TENM4 protein should be re-synthesized to reinstate function. We thus investigated the temporal dynamics of mechanosensitivity and touch sensation recovery in *Tenm4^KI/+^*mice following cutaneous PreScission injections (Figure 4A). We found that a substantial proportion of Aβ and Aδ-fibers were still mechanically silent 24h following PreScission injection in *Tenm4^KI/+^* mice. However, at 72h almost all Aβ and Aδ-fibers had mechanosensitive receptive fields, as was observed in PreScission injected WT control mice at 24 and 72h (Figure 4B). C-fiber nociceptors identified with an electrical search retained normal mechanosensitivity at all time points and conditions (Figure 4B). To assess how this recovery manifests behaviorally, we monitored paw withdrawal thresholds over time following PreScission injection. Consistent with the electrophysiological data, we found that PreScission-injected *Tenm4^KI/+^* mice displayed elevated paw withdrawal thresholds at 12, 24, and 48h which had then returned to normal levels at 72h (Figure 4C). Control WT mice injected with the PreScission protease showed no changes in paw withdrawal thresholds over the same period of time (Figure 4C). Together, these results reveal that in sensory endings the TENM4 protein is likely both necessary and sufficient for touch transduction.

**Figure 4.**
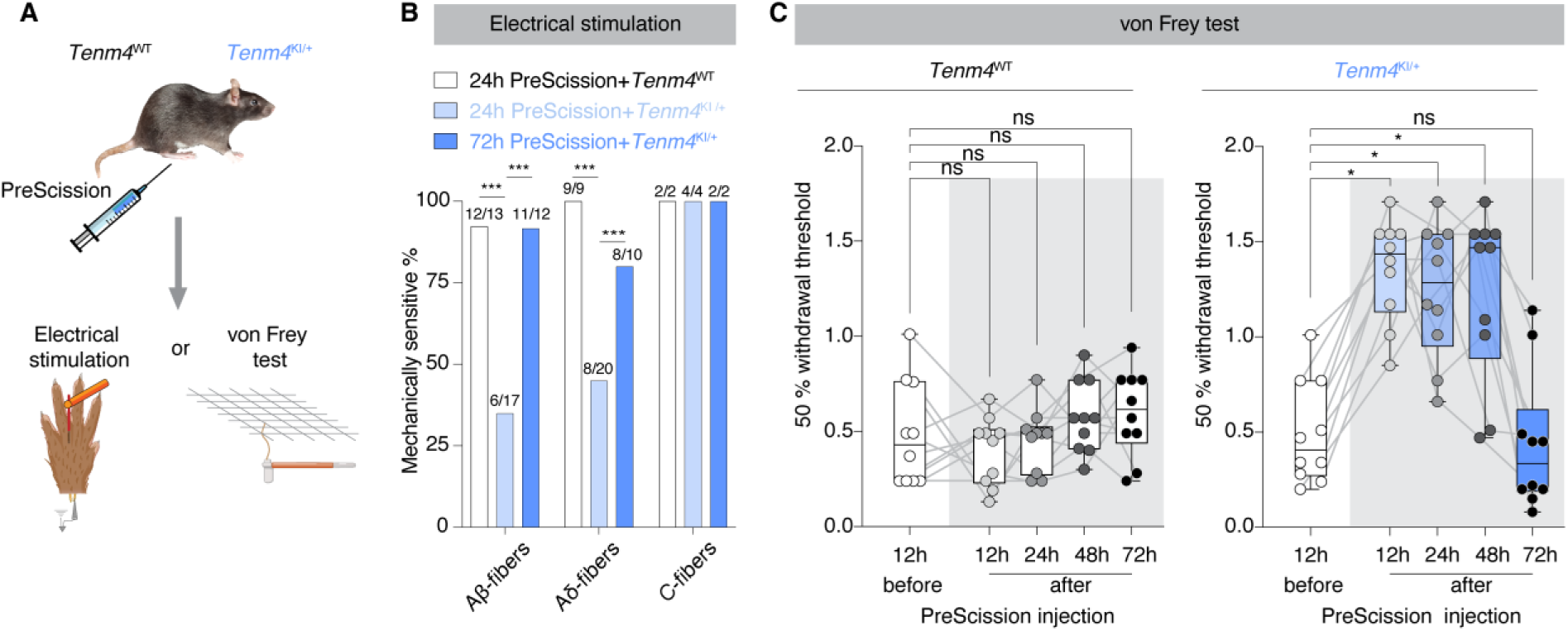
TENM4 is at sensory endings is necessary and sufficient for touch sensation. (**A**) Experimental design. (**B**) *Tenm4^KI/+^* mice exhibit a significant decrease in the proportion of Aβ and Aδ fibers displaying mechanosensitivity only after 24h, but not at 72h, following PreScission injection, while C-fibers remain unaffected at both time points. All WT fibers showed no change. Fisher’s exact test. Data were collected from 4 mice per group. (**C**) WT mice showed constant mean withdrawal thresholds to von Frey filaments before and up to 72h following PreScission treatment. In contrast, PreScission-injected *Tenm4^KI/+^* mice displayed significantly elevated paw withdrawal thresholds at 12, 24, and 48 hours, which returned to normal by 72h. Statistical significance calculated with Friedman test followed by Dunn’s multiple comparisons test. Data collected from 10 mice per group.

### TENM4 is necessary and sufficient for MA-currents

To connect behavioral recovery with underlying transduction mechanisms, we assessed mechanically activated currents in identified sensory neurons. Cultured large-diameter sensory neurons possess sensitive and large amplitude indentation-induced currents that are also sensitive to protease treatment.^12,17,47^ We made whole cell patch clamp recordings and measured MA currents in isolated large sensory neurons (diameter >30μm) from the two mouse models (*Tenm4^CKO^* and *Tenm4^KI/+^*) using cell indentation (Figures 5A and 5B). As expected almost all large neurons from control mice positive for GFP (*Tenm4^fl/fl^* injected with AAV-PHP.S-eGFP virus, lacking Cre) displayed robust MA currents (Figure 5A)^12,47^. In contrast, over half of the large diameter GFP-positive neurons recorded from *Tenm4^CKO^* mice lacked any MA current (Figure 5A). However, the incidence and characteristics of MA currents recorded from small diameter sensory neurons, which give rise to unmyelinated C-fibers, were comparable between neurons from *Tenm4^CKO^* mice and controls (Figure S8A). Similarly, a substantial proportion of PreScission-treated large-diameter DRG neurons in *Tenm4^KI/+^* showed no MA current which was significantly different from WT treated neurons (Figure 5B). Importantly, the loss of MA-currents was not accompanied by detectable changes in electrical excitability as reflected by unchanged resting membrane potentials or action potential thresholds in sensory neurons isolated from both mouse models (Figures S8B and S8C).

**Figure 5.**
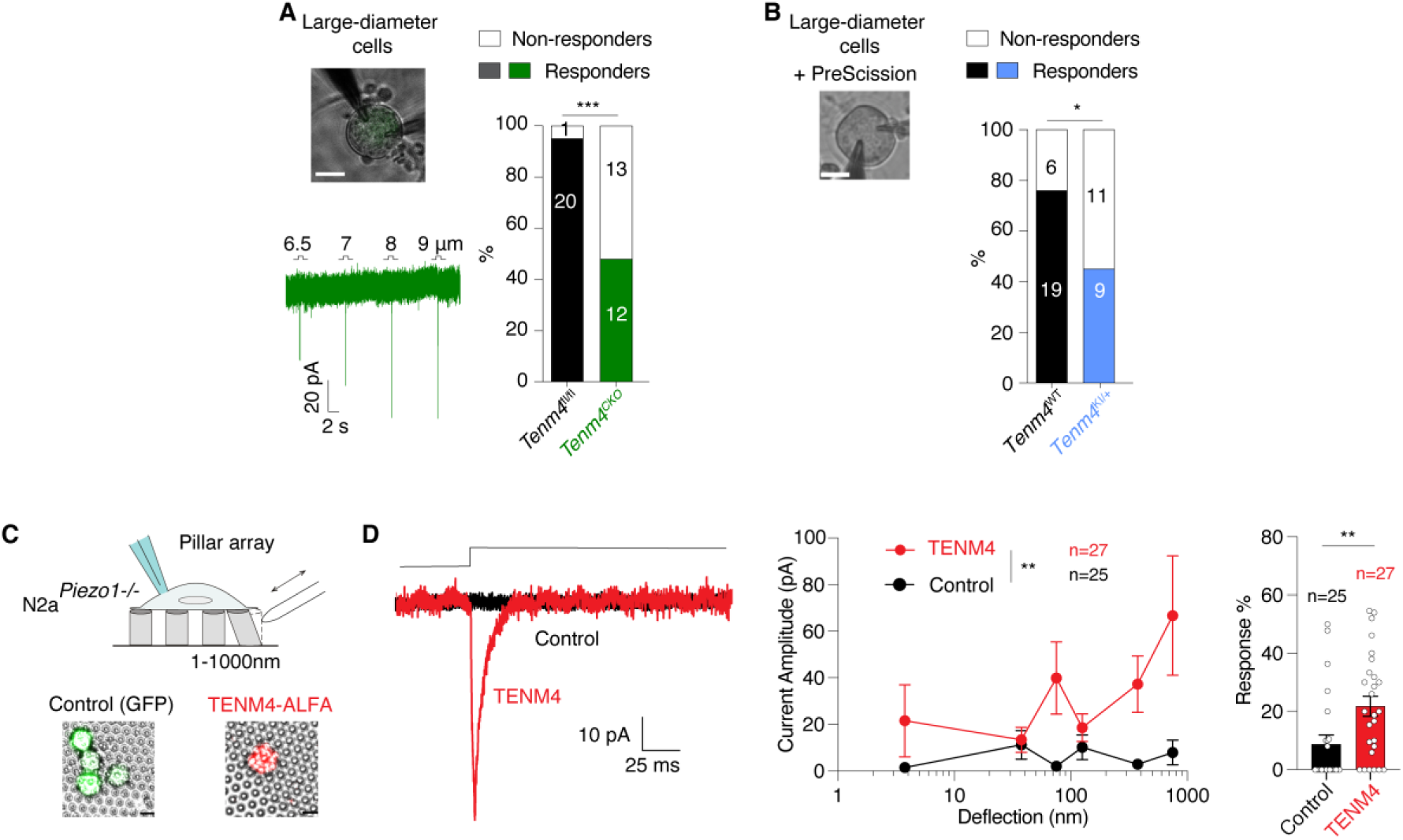
TENM4 is necessary and sufficient for MA-currents. (**A**) Poking-induced currents in GFP positive sensory neurons infected with the AAV-PHP.S-Cre-EGFP. Scale bar 20μm. *Tenm4^CKO^*mice had a deficit in MA-currents in large neurons compared to control *Tenm4^fl/fl^*mice. Significance calculated with Chi-squared test. Data collected from 4 *Tenm4^fl/fl^*and 5 *Tenm4^CKO^* mice. (**B**) Recordings from large diameter sensory in culture (scale bar = 20μm) showed a substantial decrease in the proportion of cells with poking-induced currents, specifically in cells from *Tenm4^KI/+^* treated with the PreScission protease. Significance calculated with Chi-squared test. (**C**) Cartoon of pillar stimulation. Example of transfected N2a*^Piezo1-/-^*cells expressing *Tenm4-ALFA* (red) and *pRK8-eGFP* (black) on pillar array. Scale bar 5 μm. (**D**) Representative pillar stimulation with current traces evoked from single pilus movements in N2a*^Piezo1-/-^*cells expressing *Tenm4-ALFA* (red) compared to control *pRK8-eGFP* transfected cells (black). Deflection-current amplitude plot showing large increase in the amplitude and sensitivity of MA-currents in cells transfected with TENM4 (red) compared to control (black). Statistical significance calculated with Kruskal-Wallis test. Right shows that MA-currents were evoked at much higher frequencies, 20.7± 4.2 % in *Tenm4-ALFA* expressing cells, compared to controls, 7.9 ± 3.4%. Statistical significance calculated with Mann-Whitney test.

Ion channels like PIEZO2 and ELKIN1^48,49^ are efficiently gated by substrate deflection and we asked whether TENM4 can modulate this gating.^3,12,49^ We generated full length *Tenm4* expression constructs with a C-terminal ALFA-tag that can be recognized by fluorescently labeled nanobodies (Figure S9A).^50^ We used Neuro2a-PIEZO1-Knockout (N2a*^Piezo1-/-^*) cells that have minimal, but detectable deflection induced currents,^51,52^ and transfected these cells with the *Tenm4-Alfa* or control *pRK8-eGFP* constructs. Transfected cells were plated on pillar arrays and then live labelled with anti-ALFA tag nanobodies which bind to the extracellular C-terminus of the expressed TENM4 protein. We made targeted whole cell patch clamp recordings from cells that expressed TENM4 on the membrane and measured the sensitivity and amplitude of currents evoked by single pillar displacements (Figure 5C). As observed previously, control GFP transfected cells exhibited occasional small amplitude currents to pillar deflection^51,52^. Strikingly, we observed large amplitude mechanically-gated currents in cells with plasma membrane-localized TENM4, and a quantitative analysis revealed a large increase in the incidence, amplitude and frequency of pilli-evoked currents in these cells compared to GFP controls (Figure 5D).

This data suggests that N2a*^Piezo1-/-^* cells normally express MA channels that interact with TENM4 to facilitate gating of channels via the extracellular matrix. We therefore carried out a quantitative proteomic analysis of N2a cells to reveal expressed ion channels potentially targeted by TENM4. We compared the proteome of N2a cells with that of N2a*^Piezo1-/-^*cells and could quantify the levels of more than 8000 proteins in both samples (Figures S9B and S9C). Interestingly, a large proportion of the proteome showed significant changes in the absence of PIEZO1, e.g. >16% of the proteome showed significantly higher abundances. Included in the significantly up-regulated protein set were TENM4 itself as well as the MA ion channel ELKIN1.^12,49^ We did not detect any peptides from PIEZO2, suggesting that unlike ELKIN1 this channel is not expressed in N2A cells. Interestingly, many of the highest up-regulated proteins were themselves components of the extracellular matrix.

### TENM4 associates with mechanically activated ion channels

Given the TENM4 dependence of MA currents, we examined whether TENM4 physically associates with candidate mechanosensitive channels like PIEZO2 or ELKIN1. Due to the structural similarity among Teneurins especially in the C-terminal domain, we designed a truncated construct covering the entire N-terminus and a portion of the extracellular region including sequences involved in dimerization to assess binding between Teneurin family members. Utilizing a tripartite-GFP-based protein complementation assay in HEK293T cells, we confirmed that the Teneurins can interact with each other (Figures S10A and S10B). We also measured a robust complementation signal (GFP fluorescence) between ELKIN1 and TENM4, and the strength of the interaction signal was notably much higher than that observed for ELKIN1 with other Teneurins (Figures S10C and S10D). We next asked whether TENM4 is associated with MA channels on the plasma membrane using Airyscan super-resolution microscopy. In N2A cells we co-expressed our ALFA-tagged *Tenm4* construct with either mScarlet-tagged Elkin1 or Piezo2 constructs. Nanobody-labeled TENM4 exhibited a punctate distribution on the plasma membrane as did mScarlet-tagged ELKIN1 or PIEZO2 (Figures 6A and S10E). Quantification indicated that approximately 75% of TENM4 was colocalized with ELKIN1, while around 30% of TENM4 localized together with PIEZO2 channels (Figures 6A and S10E). We next used HEK-293 cells transfected either with *Elkin1* or *Piezo2* expression plasmids and ALFA-tagged *Tenm4*. We used ALFA nanobody conjugated resins (ALFA SelectorPE^TM^) to pull down the expressed TENM4 protein complexes which were then eluted using high concentrations of the purified ALFA peptide. Using Western blotting, we could detect co-elution of ELKIN1 specifically in cases where TENM4-ALFA protein had been expressed, but not PIEZO2 (Figures 6B and S10F). Together, these results demonstrate specific and direct interactions between TENM4 and MA channels like ELKIN1. Our data does not exclude a potential interaction between PIEZO2 and TENM4 in sensory neurons as it is possible that molecular factors necessary for this interaction are not present in HEK293 or N2A cells.

**Figure 6.**
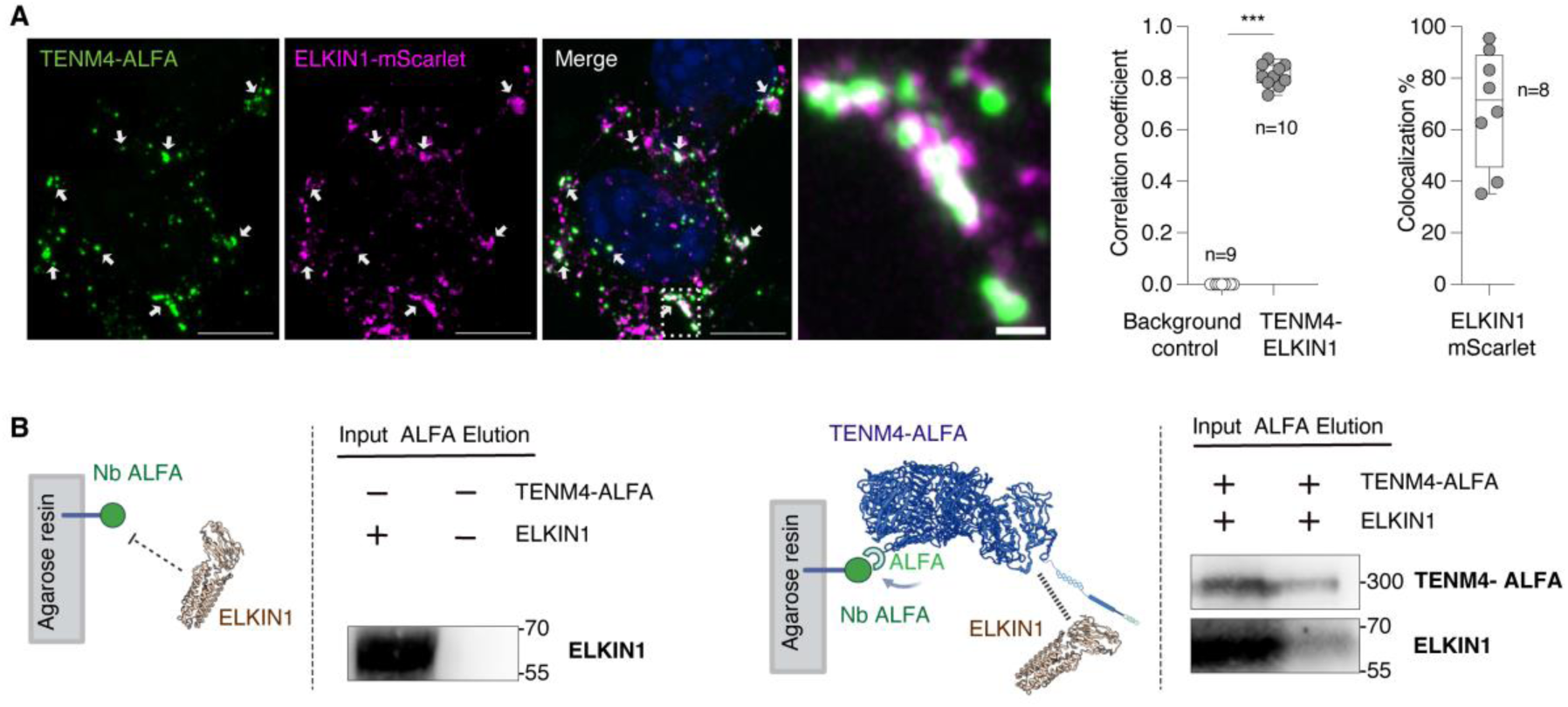
TENM4 interacts with ELKIN1. (**A**) Representative super-resolution images of N2a cells co-transfected with a *Tenm4-ALFA* expression plasmid with mScarlet-ELKIN1 (purple in middle), arrows indicate co-localization signals. Right higher magnification of co-localized protein region (from white dash boxes). Scale bar is 10μm for cells, 1μm magnified region. Quantification was carried out and a co-localization correlation, ranging from -1 to 1, signifies the strength of overlap. Values near 1 indicate strong co-localization, near -1 indicate separation. TENM4-ELKIN1 displays strong co-localization. Right, TENM4 shows 75% overlap with ELKIN1 on the cell surface. (**B**) Co-elution of TENM4-ALFA and ELKIN1 using an ALFA selector resins. Western blot of ALFA pull-down elution from HEK293T cell transfected with TENM4-ALFA and ELKIN1 (right panel) or ALFA vector plasmid and ELKIN1 (left panel). * indicates p < 0.05; ** indicates p < 0.01; *** indicates p < 0.001 means ± s.e.m.

These data suggested that the clear increase in MA currents in N2a*^Piezo1-/-^* cells overexpressing TENM4 could be due to increased interaction with endogenous ELKIN1 channels. We tested this idea by measuring deflection induced currents in N2a*^Piezo1-/-^* cells treated with siRNAs targeting the *Elkin1* transcript. Knockdown of *Elkin1* transcripts in N2a*^Piezo1-/-^* cells over expressing TENM4 did not prevent the dramatic increase in MA currents in these cells compared to controls (Figure S10F,G). This negative result may mean that TENM4 increases mechanosensitive currents in N2A cells via interaction with an as yet unknown ion channel. However, it is also possible that siRNA targeting in this case was not sufficiently efficient to remove all the ELKIN1 protein in these cells.

### TENM4 is a part of a mechanosensory tether

These molecular interactions prompted us to test a structural model in which TENM4 links mechanically activated (MA) channels to the extracellular matrix within discrete membrane domains. We have already shown that large extracellular tethers made by mouse sensory neurons in culture appear to be necessary for the expression of mechanosensitive currents.^17^ Furthermore, specific cleavage of TENM4 in mouse sensory neurons is sufficient to abolish mechanosensitive currents (Figure 5B). We therefore next asked if we could localize the protein to matrix-neurite structures in cultured mouse sensory neurons. We performed immunogold labeling with our antibody targeting the extracellular domain of TENM4 followed by transmission electron microscopy (TEM). Remarkably, immunogold labelled TENM4 was localized at the neurite-laminin interface, often associated with filamentous extracellular tether-like structures (Figures 7A, 7B, S11A, and S11B; arrowheads). Over 90% of the gold particles within 200 nm of the neurite membrane were localized to the neurite laminin interface, with only a small fraction found elsewhere, indicating strong enrichment of TENM4 at sites of extracellular contact (Figure 7C; 423 at interface vs. 45 outside). Analysis of 215 neurites revealed that while the majority of neurites harbored a single gold-labeled tether, more than 30% of neurites displayed multiple labelled tether structures in close proximity to each other on the same neurite (Figure 7B,D). This latter finding suggests that TENM4-positive tethers may often be enriched in discrete domains along the neurite (Figure 7D). Of the 325 total gold-labelled tethers analyzed, most (∼70%) were labeled with a single gold particle, however a notable fraction of labelling (∼30%) consisted of multiple gold particles on a single structure. Additionally, we calculated a density of ∼2 gold-labeled tethers/µm² of neurite laminin interface (Figures S11A and S11B), consistent with our earlier findings that contrast enhanced tether structures have a density of ∼ 2-4/µm^2^.^17^ These findings provide compelling ultrastructural evidence that TENM4 is a key component of tether-like structures at the neurite–laminin interface (Figure 7F).

**Figure 7.**
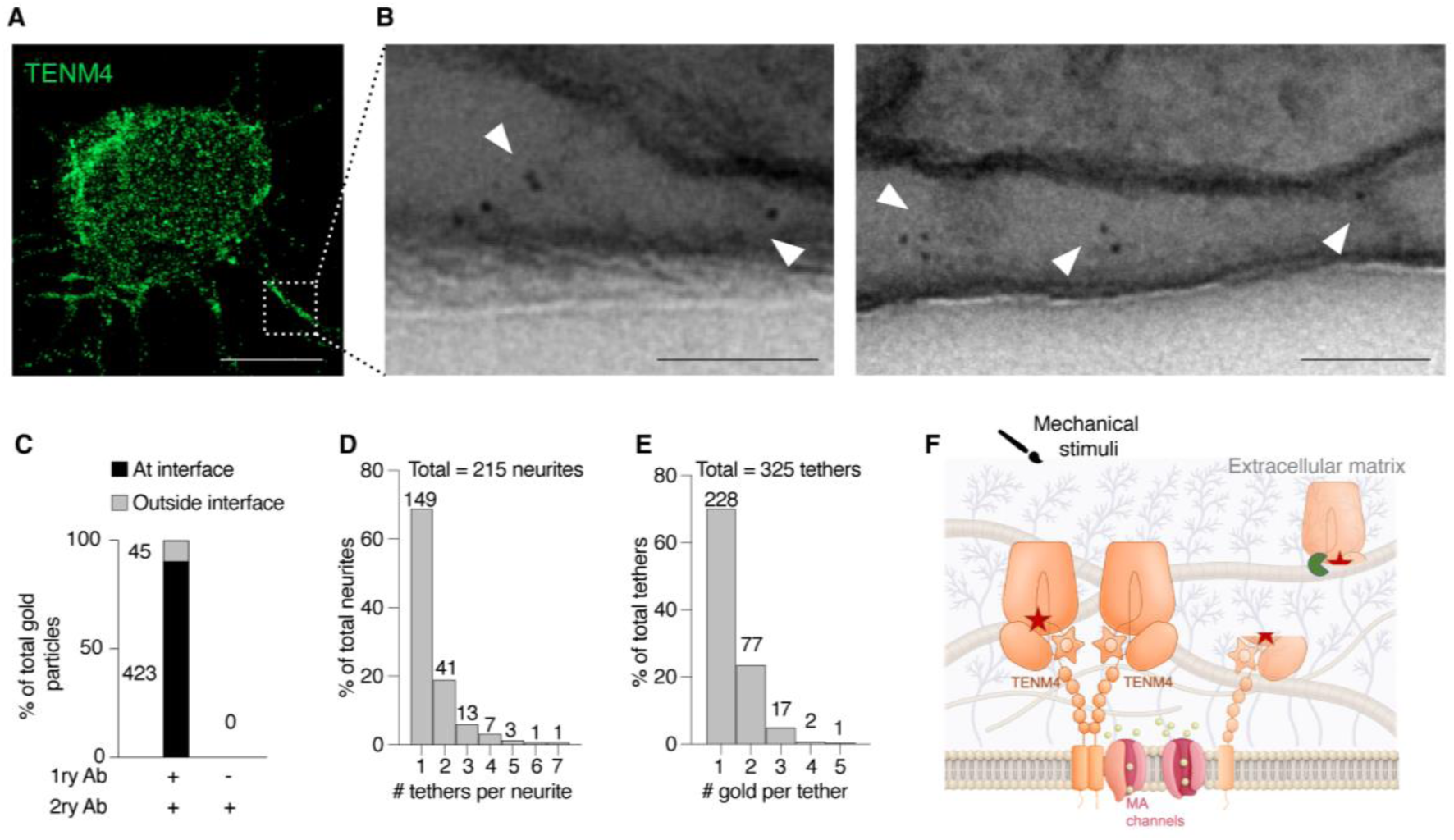
TENM4 is localized at sensory tethers. (**A**) Immunofluorescence of mouse sensory neurons stained with the custom antibody recognizing extracellular epitopes. TENM4 displays a punctate membrane pattern. Scale bar is 20μm. **(B)** Representative TEM images of two neurites showing variable Tenm4 labeling. The left neurite contains two tethers, one with a single gold particle and one with four, while the right neurite shows three tethers with one, two, and three gold particles, respectively (arrowheads) Scale bar 100 nm **(C)** Gold particles within 200 nm of the neurite membrane are highly enriched (>90%) at the neurite-laminin interface. **(D)** ∼70% of neurites have a single gold-labeled tether; ∼30% have multiple. (**E**) ∼70% of tethers carry one gold particle; ∼30% have multiple. (**F**) Cartoon illustrating how TENM4 transmits extracellular mechanical forces to ELKIN1 channels to drive mechanosensitive currents. Loss of TENM4 or its proteolytic cleavage with a protease (Green Pacman) disrupts this mechanical coupling and reduces mechanosensitive currents.

## DISCUSSION

Almost all multicellular organisms respond rapidly to the tiniest of mechanical cues. Classic genetic screens in model organisms like *Drosophila melanogaster* and *Caenorhabditis elegans* have shown that not just mechanically gated ion channels are required for mechanoreceptor function, but also a number of extracellular matrix proteins^6,9,53,54^. Body touch receptors in *C. elegans* are probably the best characterized mechanoreceptors at the molecular level and here at least a dozen proteins are required for normal mechanotransduction including three ECM proteins MEC-1, MEC-5 and MEC-9^9,53–55^. Recently, compelling evidence was provided that the Kunitz domain protein MEC-1 may link the mechanosensitive ion channel MEC-4 to specialized basal lamina at sites of transduction.^9^ Mammalian cutaneous touch receptors are diverse in terms of end-organ morphology^14^, but also in terms of the mechanosensitive ion channels required for their function. Thus in mice, genetic ablation of genes encoding the mechanically activated channels PIEZO2 or ELKIN1 or their modulator STOML3 leads to silencing of mechanoreceptor function, but in each case large sub-populations of mechanoreceptors retain normal function.^4,12,35,36^ In flies and worms genetic rescue of touch insensitivity has been routinely shown to provide evidence that the ion channel and ECM components are directly involved in transduction.^6,53,54,56–58^ Genetic rescue experiments in mouse models are more difficult, but in some cases reintroduction of mechanotransduction genes into isolated sensory neurons was shown to reconstitute mechanosensitive currents.^12,35^ In the present study we used an unbiased approach to identify a novel component of mammalian touch namely the type II membrane protein TENM4. Using sensory neuron specific *Tenm4* gene deletion in adult mice we demonstrate that presence of this protein is essential for transduction in virtually all mechanoreceptors that innervate both hairy and glabrous skin (Figure 2). In addition, genetic ablation of *Tenm4* also led to the functional silencing of fast conducting mechanonociceptors, but had no discernible effect on unmyelinated C-fiber afferents (Figure 2). These phenotypes were consistent with the relatively restricted expression of TENM4 in sensory neurons with myelinated axons (Figure 1). There were small diameter neurons positive for TENM4 e.g. small diameter tyrosine hydroxylase positive neurons (Figure 1D). However, our data on Parvalbumin positive proprioceptors suggested that not all sensory neurons are capable of transporting TENM4 to sensory terminals where transduction takes place.

TENM4 has very well characterized signaling roles within the developing CNS ^59^ and so it was important to establish that the profound sensory phenotypes we observed were not due to a failure of inter-cellular signaling at neuroglial endings. For this reason, we established a novel technology termed PreScission Protein Disassembly which relies on the use of a highly specific protease to acutely manipulate proteins in vivo with near perfect specificity. We show that acute proteolytic cleavage of TENM4 molecules at the site of transduction is sufficient to completely abolish mechanoreceptor function within minutes (Figure 3). After PreScission cleavage of TENM4, it took 72 hours for newly synthesized TENM4 to restore both mechanoreceptor function and normal touch evoked behavior. The rescue of mechanoreceptor function with endogenously expressed un-cleaved TENM4 demonstrated that the presence of intact TENM4 at the sensory ending is both necessary and sufficient for touch receptor function (Figure 4).

We hypothesized that there are protein tethers that couple mechanically gated ion channels to the ECM to facilitate fast mechanotransduction. Using cultured sensory neurons as a model we had previously shown that large tether like proteins that connect the sensory neuron membrane with the ECM appeared to be necessary for fast mechanotransduction.^17,22^ These tether-like structures were ablated by enzymes that cleave furin consensus sequences and this formed the basis of our unbiased screen which led to the identification of TENM4. Using PreScission Protein Disassembly we now show that fast mechanosensitive currents in isolated sensory neurons depend on intact TENM4 (Figure 5). This data strongly suggests that protein tethers synthesized by sensory neurons are at least in part made up of TENM4 molecules. Indeed, using immunogold TEM we show a dense localization of TENM4 at the neurite–laminin interface, where it clusters along filamentous tethers. These findings identify TENM4 as a core structural element of a putative mechanosensory tether linking the ECM to mechanosensitive channels. Interestingly, the density of TENM4 positive tethers was very similar to our previous estimates of tether density.^17,22^ Gold labelling of tethers was often found very close to the neurite membrane, consistent with the known compact structure of the extracellular teneurin domain, recognized by our antibody.^44,60^ However, we also frequently observed multiple gold labels along filaments often at distances longer than the dimensions of a single extracellular domain (∼30 nm) (Figure 7). This observation suggests that some of the labelled TENM4 may undergo regulated cleavage and then be incorporated into tethers^59^. It is intriguing in this context to note that a recent study on fly Teneurin-M suggested that this protein can form oligomeric zippers via protein interfaces.^61^ Cultured sensory neurons have served as a very useful model for mechanotransduction at the sensory ending. However, it is increasingly clear that specialized sensory Schwann cells at a variety of neuroglial endings actively participate in the transduction of mechanical stimuli.^15,62,63^ This means that examining sensory cells alone may only reveal part of the mechanistic basis of transduction.^15,64^ Here we show that TENM4 is synthesized by sensory neurons and is present at their endings in a variety of end-organ structures including Meissner’s corpuscles, Merkel cell complexes and lanceolate endings around hair follicles (Figure 1). Indeed, acute TENM4 cleavage abolishes mechanotransduction at all these sensory endings in the skin (Figure 3). It was striking that TENM4 integrity is required for transduction at such a wide variety of sensory endings suggesting that this protein is truly central to virtually all touch transduction.

One implication of our findings is that there is likely a connection between TENM4 molecules and mechanically gated ion channels. Consistently, we show that in a heterologous expression system TENM4 strongly associates with the mechanically activated ion channel ELKIN1 (Figure 6). In contrast, we were not able to co-purify PIEZO2 protein with TENM4 in the same system, but this does not necessarily mean there is no interaction in vivo. Indeed, it will be important to determine what other proteins TENM4 interacts with in sensory neurons given it’s central role in touch transduction. We also made the striking observation that introduction of TENM4 into N2a*^Piezo1-/-^* cells was sufficient to induce mechanically activated currents, however, we were not able to unequivocally determine whether this effect was exerted via ELKIN1 or by other ion channels expressed in these cells.

Teneurins play essential roles in nervous system development, in particular during neuronal migration, oligodendrocyte differentiation, and establishment of neural circuits.^44,65^ Our data provide an unexpected and surprising link between one of these proteins and fast mechanotransduction, as we show that the integrity of TENM4 is absolutely necessary for almost all somatic sensation. Indeed, the mechanisms by which Teneurins coordinate neurodevelopmental events may well include interactions with mechanosensitive ion channels the roles of which in development have only just started to be investigated.^66^ In conclusion, here we have identified a new essential component of touch transduction with a novel methodology that allowed us to definitively establish an essential role for intact TENM4 in rapid touch transduction. The central role of TENM4 for mechanotransduction in virtually all myelinated mechanoreceptors makes this molecule an ideal starting point to identifying further touch transduction components.

## ACKNOWLEDGEMENTS

We thank Maria Braunschweig, Franziska Bartelt, Kathleen Barda and Madlen Driesner for excellent technical assistance. Franziska Binder, Franziska Westphal, and Florian Keim for animal care taking. Séverine Kunz and Christina Schiel for electron microscopy and Protein Production & Characterization Technology Platform (MDC) team for excellent technical assistance. We thank members of Lewin lab for constructive discussions. We thank Stefan Lechner and Clément Verkest the mouse Piezo2-mScarlet construct.

## Author contributions

Conceptualization: M.A.K., A.T-L. H and G.R.L. Mouse model selection, design and validation: G.R.L, V.B. A.S. N.Z. and M.A.K; Patch clamp electrophysiology: S.C (poking) and J.A.G-C. (Pillar array); Antibody validation: MAK, L.D., R.G. A.S. and A.T-L.H; Tripartite GFP design and implementation; candidate screening and molecular biology: YASB, L.D and A.T-L. H. Skin nerve electrophysiology: M.A.K, R.G, W.Z, S.J.M. and G.R.L; Super resolution microscopy and biochemistry: A.T-L.H, M.A.K and S.H.; immunogold labelling-TEM: L.D., S.K. C.S., and E.N.; Behavioral assessments M.A.K. with R.G. Protein Production: A.S. Proteomics J.A.G-C, S.C, O,P and P.M. Writing: M.A.K. and G.R.L with input from all authors; Supervision and funding: G.R.L.

## SUPPLEMENTAL INFORMATION

Method details

Figures S1 to S11

Tables S1 to S2

