## Supplementary Methods and Figures for "TENM4 is an essential transduction component for touch"

**The file includes:**

Method details

Figures S1 to S11

Tables S1 to S2

References

**Methods details**

**Animals**

All experiments were approved by the Berlin Animal Ethics Committees (Landesamt für Gesundheit und Soziales) and carried out in accordance with European animal welfare law. Wild-type (C57BL/6N) adult (8-30 weeks) mice of both sexes were used. Mice were housed in groups of 5 with food, water and enrichment available ad libitum.

The *Tenm4*$^{KI/+}$ mouse line was generated by the Ingenious Targeting Laboratory (Ronkonkoma, NY) using homologous recombination in mouse ES cells. Targeted iTL IN2 (C57BL/6) embryonic stem cells were microinjected into Balb/c blastocysts. Resulting chimeras with a high percentage black coat color were mated to C57BL/6 FLP mice to generate Somatic Neo Deleted mice. This involved the replacement of the amino acids HLYTQSL (exon 27) with the PreScission Protease target sequence LEVLFQG, as illustrated in Figure S2. After the removal of the Neo cassette, a single FRT site of 78 bp is retained and subsequently utilized for the selection process and genotyping. The genotyping strategy utilized was as follows: Forward primer: 5'- GGGAAATGGGATGGTGGGAA -3', reverse primer: 5'- ACCCACGTGGAGTTGTTCTTC -3'. The expected band for WT is 231 bp, while for *Tenm4*$^{KI/+}$, two bands are observed at 231 and 309 bp.

*Frem2*$^{fl/fl}$ and *Tenm4*$^{fl/fl}$ mouse lines were generated at Shanghai Model Organisms Center using the CRISPR/Cas9 strategy. *Frem2*$^{fl/fl}$ *and Tenm4*$^{fl/fl}$ mice, carring LoxP sites flanking exon 12 and exon 7, respectively, were generated using the following gRNAs *Frem2*$^{fl/fl}$-gRNA1 (5'-ACAAGACTAGCTCTGGGTCTGGG -3'), *Frem2*$^{fl/fl}$-gRNA2 (5'-TACAGAGATAACAGTGGGGGTGG -3'), *Tenm4*$^{fl/fl}$-gRNA1 (5'-AGAGCCCAGTCTCTCACCAGTGG -3') and *Tenm4*$^{fl/fl}$-gRNA2 (5'-GCACTGAACCTCAGCCCTGGTGG -3'). The *Frem2*$^{fl/fl}$ genotyping strategy utilized was as follows: forward primer (5'- AGTCGGGTTTCTTCTTCCACAA -3'), reverse primer (5'-CCCAAAAGGTCACATGCTAATCTAT -3'). The expected band for WT is 197 bp, while for *Frem2*$^{fl/fl}$, a band with 263 bp. The *Tenm4*$^{fl/fl}$ genotyping strategy utilized was as follows: forward primer (5'-TGAACCATGCATCCCTCACC -3'), reverse primer (5'- AAACTCCCACCCCACTGAAC -3'). The expected band for WT is 313 bp, while for *Tenm4*$^{fl/fl}$, a band with 379 bp. Animals of various genotypes were randomly selected for experiments based on their availability.

*Pv-Cre:Tenm4*$^{fl/fl}$ mice were generated by crossing Pv$^{Cre}$ (B6;129P2-*Pvalb*$^{tm1(cre)Arbr}$/J; RRID:IMSR_JAX:008069) mice and *Tenm4*$^{fl/fl}$ mice. *PV*$^{cre}$;*Rx3*$^{flpo}$;Ai65D mice were generated by crossing *PV*$^{cre}$ and *Rx3*$^{flpo}$ lines with the Ai65D reporter allele to create a triple transgenic line that selectively labels proprioceptive sensory neurons.[1–3] Muscle spindles and Golgi tendon organs (GTOs) are unique in that they are the only cells in the mouse body that co-express parvalbumin (PV) and the transcription factor RUNX3 (Rx3). This molecular intersection provides a highly specific genetic entry point for targeting proprioceptors.[3] In the *PV*$^{cre}$;*Rx3*$^{flpo}$;Ai65D line, cre- and flp-dependent recombination at the Ai65D locus induces robust expression of tdTomato exclusively in proprioceptive DRG neurons and their peripheral and central projections. This strategy enables the precise anatomical visualization of proprioceptive terminals within muscle and provides a reliable reporter for experiments requiring the identification of spindle and tendon organ afferents.

**AAV transduction**

*pAAV-Syn-Cre-p2A-EGFP* (4.91 x $10^{12}$ vg/ml) and *pAAV-hSyn-NLS-GFP-P2A* (4.98 x $10^{12}$ vg/ml) were manufactured in the Charité Viral Core facility (Berlin, Germany). *Frem2*$^{CKO}$ 765 and *Tenm4*$^{CKO}$ were generated by retroorbital injection of Adeno-associated virus (*pAAV-Syn-Cre-p2A-EGFP*) in *Frem2*$^{fl/fl}$ and *Tenm4*$^{fl/fl}$, respectively, to induce Cre/GFP recombinase expression in DRG neurons. Subsequently, DNA between the LoxP sites will be excised, leading to a frameshift mutation and the production of a truncated protein as illustrated in Figure S2. Control mice were generated by similar injections of *pAAV-hSyn-NLS-GFP-P2A* that induce GFP expression in DRG neurons. Afterwards the animals were returned to their home cage for at least three weeks to achieve an adequate recombination and subsequently tested in electrophysiological and behavioral assays.

**Tamoxifen injections**

Tamoxifen-induced activation of Cre recombinase under the Advillin promoter in sensory neurons was used as a second inducible approach to delete *Tenm4* using the AdvillinCreERT2 transgenic mice (*Tenm4*$^{fl/fl}$*; Avil*$^{CreERT2/WT}$). Briefly, *Tenm4*$^{fl/fl}$*; Avil*$^{CreERT2/WT}$ mice were injected intraperitoneally with corn oil (control animals) or Tamoxifen (15mg/ml) solubilized in corn oil (Sigma) for 5 consecutive days. Prior to injection, each mouse was weighed to ensure the administration of a uniform tamoxifen dose (150 mg/kg). Behavioral assays were performed 21 days after vehicle or tamoxifen injections.

**PreScission injections and ex vivo cleavage assay**

Adult mice (8-30 weeks old) were anesthetized using a 5% isoflurane-air mixture in a transparent box, with anesthesia maintenance via a 2.5% isoflurane-air mixture through a plastic tube. After achieving slow, steady breathing, a hind paw pinch test assessed pain reflexes. Subsequently, mice received a 10µL injection of saline NaCl solution (control group) or a 20% diluted PreScission protease in saline NaCl solution (v/v) 10 U/µL, Cytiva, Lot: 18043250). The injection was directed into the intraplanar surface of glabrous skin or the top of the hairy skin of the hind paw. The mice were then used for for immunohistochemistry, electrophysiological, or behavioral experiments.

For *ex vivo* PreScission protein cleavage assay, *Tenm4*$^{KI/+}$and WT mice were administered injections of 20% PreScission protease (2-10 U/µL, Cytiva, Lot: 18043250) diluted in NaCl (v/v) in the glabrous skin of hind paws. Tissues were collected one-hour post-injection for protein extraction. Frozen tissues (-80$^{o}$C) were powdered using a mortar and added to lysis buffer (RIPA) containing 50 mM Tris pH 7.5, 150 mM NaCl, 0.1% SDS, 1.0% NP-40, 0.5% deoxycholate, and a protease inhibitor cocktail tablet. Sonication (magnitude 3) with 10 seconds on and 50 seconds off was applied three times to each sample. The lysate was centrifuged at 13,200xg for 30 minutes. Samples corresponding to 1/100 of the input were resolved by SDS–PAGE. Analysis was conducted by Western blotting using a polyclonal sheep-raised antibody against TENM4 (dilution 1:500), followed by an HRP-conjugated goat anti-sheep IgG (R&D Lot: 1522051, dilution 1:10,000). Blots were developed using the SuperSignal™ West Dura ECL Kit (Thermo Scientific) and imaged using a Bio-Rad ChemiDoc™ MP imaging system.

**Plasmids and cloning**

Teneurins 1-4 and Elkin1 were independently cloned into modified plasmids, incorporating the BsmBI restriction site as detailed in the tripartite-GFP method section. This involved using specific primers for each construct: TENM1 (1-526): forward primer (5'-cacgtctcactccatggagcaaacagactgcaaa -3'), reverse primer (5'-cacgtctcacttattcaattgctgtagttagcacaaat-3'), TENM2 (1-571): forward primer (5'-cacgtctcacttattcaattgctgtagttagcacaaat -3'), reverse primer (5' ggatctgaattcttaatctaagacaacagtgttgaaggag -3'), TENM3 (1-513): forward primer (5'-cacgtctcactccatggatgtgaaggaacgcag -3'), reverse primer (5'-cacgtctcacttactctataacgatcgtgttaaaggagac -3'), TENM4 (1-561): forward primer (5'-cacgtctcactccatggagccagaccactcg -3'), reverse primer (5'- cacgtctcacttaatccagatactggatgaaaccgg -3'), and Elkin1 (full length): forward primer (5'- cacgtctcacaccatggcggtggctgc -3'), reverse primer (5'-cacgtctcatcccctccatctttgacctttcaaagtg 3') (Clontech). The full-length Tenm4 cDNA, sourced from VectorBuilder.inc (Mouse MGC Verified FL clone, ID: NM_011858.4, VB221104-1135), containing a modified C-terminus ALFA tag. Additionally, the full-length PIEZO2 cDNA with an mScarlet red fluorescent 820 protein tag plasmid was obtained as a gift from Professor Stefan Lechner at UKE Hamburg. Furthermore, the recently-published full-length ELKIN1 pRK8-eGFP,[4] was adapted to incorporate mScarlet at the N-terminus using specific primers forward primer (5'-ctatcgattgaattaagcttatggtgagcaagggcgag -3'), reverse primer (5'-caccgccatggtggcggatcccttgtacagctcgtccatgcc -3') (Clontech).

**Cell line culture**

HEK293T cells were cultured in DMEM-Glutamax (Gibco, ThermoFisher Scientific) supplemented with 10% fetal bovine serum (FBS, PAN Biotech GmbH) and 1% penicillin and streptomycin (P/S, Sigma-Aldrich). N2a cells were cultured in DMEM-Glutamax (Gibco, ThermoFisher Scientific) supplemented with 45% Opti-MEM (Gibco, ThermoFisher Scientific), 10% FBS, and 1% penicillin and streptomycin (P/S, Sigma-Aldrich). For downstream analyses, HEK293T and N2a cells were transiently transfected by plating them on glass coverslips coated with poly-L-lysine and laminin at various cell densities and maintained overnight in a 37°C, 5% CO2 incubator. Transfections were carried out using 4µL FuGene HD per 50µL of Opti-MEM with 1µg of cDNA, following the manufacturer's instructions. Subsequent analyses were performed 48 hours post-transfection, and data were derived from a minimum of 3 independent transfections.

**siRNA transfection**

2 hours after plating the N2a cell in glass bottomed dishes coated with PLL and laminin, siRNA transfection was carried out using DharmaFECT (Horizon) reagents according to the manufacturers' guidelines. Briefly, siRNAs were mixed with serum and antibiotics free plating medium in a tube (total volume of 100 µl per dish), and 4.5 µl of DharmaFECT 1 transfection reagent was mixed with serum and antibiotics free medium (total volume of 100 ul) in another tube. Each tube was incubated separately for 5 min at RT, then mixed together and incubated for a further 20 min. 800 µl of antibiotics free complete medium was added to this mixture and added to the cells. Experimental dishes were transfected with a final concentration of 50 nM ON-TARGETplus SMARTpool mTmem87a and 50 nM of siGLO green transfection indicator and control dishes were transfected with 50 nM of ON-TARGETplus Non-targeting Pool and 50 nM of siGLO green transfection indicator. Cells were transfected with mTenm4 using FuGene HD 24 hours after siRNA transfection and patched 48 hours post-siRNA treatment. n=3.

**Immunocytochemistry**

Cells were fixed in 4% paraformaldehyde in PBS for 10 minutes at room temperature, followed by three washes with PBS. Subsequently, cells were blocked for 1 hour with 10% horse serum in PBS, either in the presence or absence of 0.1% (v/v) Triton X-100 for staining intracellular or extracellular epitopes, respectively. Following blocking, cells were incubated overnight at 4°C with primary antibodies, appropriately diluted in either 3% horse serum in PBS or 3% horse serum in 0.1% (v/v) Triton X-100/PBS, depending on the location of the epitope, for epifluorescence staining or cellular staining, respectively. Primary antibodies included polyclonal sheep anti-TENM4 antibody at a dilution of 1:500 (R&D Systems (TENM4-N), Catalog #: AF6320), our own polyclonal rabbit anti-TENM4 antibody at a dilution of 1:1000 (TENM4-C) (Eurogentec), and FluoTag®-X2 anti-ALFA at a dilution of 1:500 (NanoTag, N1502-Ab635P-L). When FluoTag®-X2 anti-ALFA was employed for Alfa-tagged protein detection, the incubation time was reduced to 1.5 hours at room temperature. Following three washes with PBS, cells were incubated for 1.5 hours at room temperature with species-specific conjugated secondary antibodies diluted at 1:500 in PBS: Donkey anti-Sheep IgG (H+L) Cross-Adsorbed Secondary Antibody, Alexa Fluor™ 568 (Invitrogen, Catalog # A-21099), and goat anti-Rabbit IgG (H+L) Cross-Adsorbed Secondary Antibody, Alexa Fluor™ 555 (Invitrogen, Catalog # A-21428). This step was omitted when using FluoTag®-X2 anti-ALFA. After three washes in PBS, cells were stained with DAPI diluted 1:1000 in PBS for 10 minutes at room temperature, followed by three additional washes in PBS, and finally, mounting with Dako (Agilent, S3023).

**Multiplex fluorescent in situ hybridization**

*Tenm4*, *Kcnq4*, *Elkin1* and *Piezo2* mRNA levels were assessed in DRGs from WT (n = 4) mice using RNAscope Multiplex Fluorescent Kit v2 (a fluorescent *in situ* hybridization technology). Briefly, the 12μm DRG sections were immersed in PBS for 15 minutes and dehydrated in a series of 50 %, 70 %, 100 % ethanol for 5 minutes each. Afterwards, sections were treated with a hydrogen peroxide solution for 15 minutes at room temperature to block endogenous peroxidase activity followed by another wash with 100% ethanol for 5 minutes. Next, Protease plus was applied for 30 minutes at RT. After three washes with PBS, probes were applied, and hybridization was carried out in a humified oven at 40°C for 2h. Following hybridization, amplification was performed using Amp1, Amp2, and Amp3 each for 30 minutes at 40°C. For detection, each section was treated sequentially with channel specific HRP (HRP-C1, HRP-C1, HRP-C3) for 15 minutes, followed by TSA-mediated fluorophore binding for 30 minutes and final HRP blocking for 15 minutes (all steps at 40°C). The following probes were used: *Tenm4*- Mm-*Tenm4* (C1 #555491), *Kcnq4*- Mm-*Kcnq4* (C2 #472271), *Piezo2* - Mm-*Piezo2*-E43-E45 15 (C1 #439971) and *Elkin1* - Mm-*Tmem87a* (C3 #868581). The fluorophores applied to detect the signal from the probes were Opal 690 Reagent (1:750; Akoya Biosciences, #FP1497001KT) or Opal 570 Reagent (1:750; Akoya Biosciences, #FP1488001KT). Slides were rinsed with PBS-tween, followed by a 1-hour incubation at room temperature in an antibody diluent solution. Following this, they were treated for antibody immunostaining according to procedures outlined below. Lastly, the slides were exposed to RNAscope DAPI during the final incubation. Images were obtained using Airyscan fluorescence microscope (Axio, Zeiss). Exposure levels were kept constant for each slide and the same contrast enhancements were made to all slides. In the quantification experiments, over 700 neurons were counted in each group, with a representation of 2 male and 3 female mice in each group.

**Generation of anti TENM4 antibody (Anti-TENM4-C)**

The antigen used for the generation of this polyclonal antibody consisted of most of the extracellular domain of murine TENM4 (UniProt ID Q3UHK6), including the large YD shell. The expression construct for the production of the antigen was synthesized by VectorBuilder GmbH (pRP[Exp]-CMV) and comprised amino acids 921 to 2771, resulting in secretion,[5] (μphosphatase secretion leader sequence MGILPSPGMPALLSLVSLLSVLLMGCVA*ETG) of a C-terminal His6-tagged protein. The protein was produced in ExpiCHO-S™ cells (Thermo Fisher Scientific) at 31°C and 8% CO2 using BalanCD transfectory CHO medium (FUJIFILM Irvine Scientific) supplemented with 4 mM GlutaMAX (Thermo Fisher Scientific). Cells were transiently transfected using PEI MAX (Polysciences) and supplemented with 6 g/L glucose (Sigma) three days after transfection. Cell culture supernatant was collected six days after transfection, filtered through 0.22 µm filters, and supplemented with 0.5 M NaCl, 5 mM imidazole, and 25 mM sodium phosphate buffer pH 7.8. The protein was captured from the supernatant by affinity chromatography using cOmplete™ His-Tag Purification Resin (Roche), equilibrated with 50 mM sodium phosphate pH 7.8, 0.5 M NaCl, and 5 mM imidazole. The bound protein was first washed with 50 mM sodium phosphate pH 7.8, 0.5 M NaCl, followed by a buffer containing 10 mM imidazole to remove contaminating proteins. The protein was eluted with the same buffer containing 0.25 M imidazole. The protein was further purified by gel filtration on a 16/60 HiPrep Sephacryl S-400 HR column (Cytiva) equilibrated with PBS buffer pH 7.4 and 0.2 M NaCl. The purified protein was concentrated to about 3 mg/mL, sterile filtered, flash-frozen in small aliquots with liquid nitrogen and stored at -70°C until further use.

The purified antigen was then used by Eurogentec for immunization of rabbits. Two animals were injected 4 times with 100 µg of antigen. The final bleed was collected 28 days after the first injection and subsequently concentrated using affinity purification.

**Immunohistochemistry**

Mice were euthanized by cervical dislocation, and tissues including hairy and glabrous skin, sciatic nerves, and DRGs, were collected. The hairy skins were shaved, and the hypodermis, ligaments, and attached muscle tissue were removed from the skin samples. Subsequently, skin samples were stretched out using insect pins and fixed in 4% paraformaldehyde (PFA) for a duration of 4 hours then post-fixed at 4 °C for 24 h in 20% dimethylsulfoxide and 80% methanol, while sciatic nerve and DRG samples were fixed in 4% PFA for a duration of 2 hours. L2–L5 DRG and sciatic nerves were dissected and postfixed in PFA for 1 hour. Samples underwent cryoprotection with an overnight incubation in 30% (w/v) sucrose solution in 5% PBS at 4 °C, followed by embedding in optimal cutting temperature compound (OCT compound; Thermo Fisher Scientific).

$PV^{cre}$;$Rx3^{flpo}$;Ai65D mice were deeply anaesthetized and transcardially perfused with 1xPBS, followed by 4% PFA in 1xPBS. The *tibialis anterior* muscles were then dissected and post-fixed in 4% PFA at 4 °C overnight, and subsequently cryoprotected in 30% sucrose in 1xPBS until they sank. The muscles were then embedded in OCT.

Abdominal human skin samples were obtained from patients that underwent plastic surgery (written consent obtained, ethics approval from Charité University Hospital EA1/356/21). Following surgical removal, excessive subcutaneous fat tissue was removed and skin sheets were either directly used or stored at -20 °C until further usage.

OCT-embedded samples were snap-frozen on dry ice and stored at −80 °C. Sciatic nerve- and DRGs-embedded tissues were sectioned at 12 µm using a Leica Cryostat (CM3000; Nussloch), mounted on

Superfrost-Plus microscope slides (Thermo Fisher Scientific) and stored at −80 °C until staining. Skin-embedded tissues were sectioned at 60 µm and stored in freezing solution (30% (v:v) glycerol, 30% ethylene glycol, and 40% PBS). During staining, slides and free-floating skin sections were washed with PBS-tween and blocked in antibody diluent solution containing 0.2% (v/v) Triton X-100, 5% (v/v) horse serum, and 1% (v/v) bovine serum albumin in PBS for 1 hour at room temperature. Subsequently, slides underwent a 12-hour incubation, while skin samples were incubated for 72 hours at 4 °C with primary antibodies. OCT-embedded muscle tissue was sectioned at 50 µm using a cryostat (Leica CM3050 S; Nussloch), mounted on Superfrost-Plus microscope slides (Thermo Fisher Scientific) and stored at −80 °C until staining. The following primary antibodies were used: anti-TENM4 (host: sheep, AF6320 and ab215052, 1:500), anti-S100 (host: rabbit, 15146-1-AP, 1:1000), anti-NF200 (host: chicken, AB72996, 1:1000), anti-CK20 (host: guinea pig, BP5080, 1:500), anti-CGRP (host: rabbit, 24112, 1:500), anti-TH (host: sheep, AB1542, 1:1000), anti-TRKC (host: goat, AF1404, 1:1000), anti-Frem2 (host: rabbit, sc-98471, 1:400), anti-Slit1 (host: rabbit, ab10984, 1:400), anti-Scribble (host: rabbit, ab154067, 1:400), anti-Pcsk5 (host: rabbit, TA332044, 1:400), anti-Tecta (host: mouse, ABNOH00007007-A01, 1:100). Slides and free-floating skin sections were then washed three times using PBS-tween and incubated with the following species-specific conjugated secondary antibodies at 1:500 dilution: Alexa Fluor 488 anti-rabbit (A21206), Alexa Fluor 568 anti-sheep (A21099), Alexa Fluor 568 anti-chicken (A11041), Alexa Fluor 555 anti-goat (A21432), Alexa Fluor 568 anti-guinea pig (106-165-003), or Isolectin GS B4 conjugated to Alexa 594 (121413, Thermo Fisher) overnight at 4 °C. The secondary antibody was washed three times in PBS tween. DRG and sciatic nerve slides were then mounted with Dako (Agilent, S3023), muscle slides were mounted with Mowiol Mounting Medium (Roth, 0713.2), while skin samples were processed for tissue clearing using 2,2'-thiodiethanol (TDE, Sigma-Aldrich). Skin sections were then placed in increasing concentrations of TDE every 2 h, from 10% to 25%, 50% and 97%, in which samples were stored and mounted on to slides and cover slipped. Slides and free-floating skin sections were imaged with Airyscan fluorescence microscope (Axio, Zeiss). Muscle slides were imaged with a confocal laser scanning microscope (Zeiss LSM 800). Exposure levels were kept constant for each slide and the same contrast enhancements were made to all slides. Negative controls without the primary antibody showed no staining with either secondary. In the quantification experiments, over 700 neurons were counted in each group, with a representation of 2 male and 3 female mice in each group.

**DRG neuron culture**

Lumbar DRGs (L1-L6) were collected from mice into plating medium (DMEM-F12, Gibco, ThermoFisher Scientific) supplemented with 10% fetal horse serum (FHS, Life Technologies) and 1% penicillin and streptomycin (P/S, Sigma-Aldrich). A single-cell suspension was obtained by enzymatic digestion, first in 1.25% Collagenase IV (1 mg/ml, Sigma-Aldrich) for 1 hour at 37°C and then in 2.5% Trypsin (Sigma-

Aldrich) for 15 minutes at 37°C, followed by mechanical trituration with a P1000 pipette tip and purification on a 15% fraction V BSA column. Subsequently, neurons were plated on glass-bottomed dishes coated with poly-L-lysine and laminin for indentation assays or immunostaining. The cultured neurons were incubated in a standard incubator (37°C and 5% CO2). Electrophysiological experiments involving indentation were conducted 18-24 hours after plating, while immunostaining experiments were performed 48 hours after plating. For the MA-current analysis of $\textit{Tenm4}^{fl/fl}$ animals, n = 4 mice underwent AAV-GFP injection (control), and n = 5 mice underwent AAV-Cre injection. For the MA-current analysis of the effects of PreScission on WT and $\textit{TENM4}^{KI/+}$ mice, n = 5 WT, and n = 4 $\textit{TENM4}^{KI/+}$ mice were utilized. Sample sizes were determined based on previous studies with similar effect sizes.

**Real-time PCR**

Total RNA was extracted from the skin, DRG, and sciatic nerve of WT mice using the ReliaPrep™ RNA Miniprep Systems and retro-transcribed into cDNA using the GoScript™ Reverse Transcriptase kit, according to the manufacturer's instructions. The expression of TENM4 was evaluated using SYBR Green Real-Time PCR (Thermo Fisher Scientific™). The PCR mix contained 2μl of a 1:10 diluted cDNA template; 5μl of 2x SYBR® Green Master Mix (BioRad); 0.3μl of a 10μM forward primer; 0.3μl of a 10μM reverse primer; 0.5μl of DMSO; 1.9μl of H2O, in a total volume of 10μl per sample. The primers used were the forward primer (5' – cccatcagcaactctcaggac -3') and the reverse primer (5'-ccccgaggatagacttgct -3'). Standard reactions were carried out using a CFX Connect RealTime PCR Detection System. Technical triplicates were included in each experiment. Standard curves obtained with the amplification of standard plasmids at increasing known concentrations were used to quantify TENM4 expression in different tissues.

**Whole-cell patch clamp**

Neurons or transfected $N2a^{Piezo1-/-}$ cells [6,7] were immersed in an extracellular solution composed of (in mM): 140 NaCl, 4 KCl, 2 CaCl2, 1 MgCl2, 4 Glucose and 10 HEPES, pH adjusted to 7.4 with NaOH. For the MA-current analysis of the effects of PreScission on WT and $\textit{Tenm4}^{KI/+}$ mice, neurons were immersed in the identical extracellular solution, supplemented with PreScission protease at a ratio of 2:1000 (v/v), 30 minutes prior to recordings. Patch pipettes (3-6 MΩ resistance) were fashioned from heat-polished borosilicate glass (Harvard apparatus, 1.17 mm x 0.87 mm) and filled with an intracellular solution containing (in mM): 110 KCl, 10 NaCl, 1 MgCl2, 1 EGTA, and 10 HEPES, with pH adjusted to 7.3 using KOH. Current-clamp experiments involved injecting steps of current in 50 pA increments from 0-950 pA to categorize sensory neurons into mechano- (large neurons) and nociceptors (smallmedium neurons). Currents from neurons or transfected $N2a^{Piezo1-/-}$ cells were evoked using pillar deflection at a holding potential of -60 mV. Recordings were acquired with an EPC-10 amplifier with Patchmaster software

(HEKA, Elektronik GmbH, Germany). For indentation experiments, neuronal cell membrane indentation in the range of 1 – 9 µm was achieved using a heat-polished borosilicate glass pipette (mechanical stimulator) manipulated by a MM3A micromanipulator. Bright field images were captured with a 40X objective and a CoolSnapEZ camera (Photometrics, Tucson, AZ) on a Zeiss 200 inverted microscope for the analysis of neuronal diameter. Currents and biophysical parameters were analyzed using FitMaster (HEKA, Elektronik GmbH, Germany).

**Pillar arrays**

Pillar arrays were prepared as described previously.[4,6,8,9] Briefly, salinized negative masters were used as templates. Negative masters were covered with degassed polydimethylsilozane (PDMS, syligard 184 silicone elastomer kit, Dow Corning Corporation) mixed with a curing agent at 10:1 ratio. After 30 min, glass coverslips were placed on the top of the negative masters containing 35 µm PDMS mix and baked at 110˚C for 60 min. The pillar arrays were peeled away from the master. The resulting dimensions of single pilus within the array was radius= 1.79 µm; length= 5.87 µm. While the elasticity was 2.1 MPa and the spring constant was 251 pN-nm as previously reported.[4,6,8,9] The pillar arrays were plasma cleaned (Deiner Electronic GmbH, Germany) and coated with PLL, subsequently N2a$^{Piezo1-/-}$ cells were cultured and were prepared as previously described. 24 hours after cells adherent on pillar, cells were transfected with either pRP TENM4-ALFA or pRK8 eGFP. After 24 hours of transfection, all cells were stained with FluoTag®-X2 anti-ALFA, as described earlier without fixation and Triton X-100. GFP cells or live-staining positive cells from respective cultures were selected for subsequent experiment.

To create quantitative data an individual pilus subjacent to the cell membrane was deflected using a heat-polished borosilicate glass pipette (approx. 2 mm in diameter) driven by a MM3A micromanipulator (Kleindiek Nanotechnik, Germany) as previously described in.[4,6,8,9] Deflection stimuli were applied to a single pilus in the range of 1-1000 nm and cells were monitored using whole-cell patch-clamp. A bright-field image (Zeiss Axio Observer A1 inverted microscope) was taken before and during pillar deflection stimuli using a CoolSnapEZ camera (Photometrics, Tucson, AZ) and 40x objective. The pillar deflection was calculated by comparing the light intensity of the center of each pilus before and after the stimuli with a 2D-Gaussian fit (Igor Software, WaveMetrics, USA). Stimuli larger than 1000 nm were excluded.

**Ex vivo skin nerve**

Ex vivo skin nerve electrophysiology was performed on cutaneous sensory fibers of the tibial and saphenous nerves, following a previously established method.[10] Briefly, mice were euthanized by cervical dislocation, and the hair on the limb was shaved off. The hairy skin from the hind paw, along with the saphenous nerve up to the hip, was then dissected. The innervated hairy skin was placed in a bath chamber continuously perfused with warm (32°C) carbonated (95% O2, 5% CO2) interstitial fluid (SIF buffer): 123

mM NaCl, 3.5 mM KCl, 0.7 mM MgSO4, 1.7 mM NaH2PO4, 2.0 mM CaCl2, 9.5 mM sodium gluconate, 5.5 mM glucose, 7.5 mM sucrose, and 10 mM HEPES (pH 7.4). The skin was stretched and fixed "inside-out" using insect needles, allowing stimulation of the inner part of the skin with stimulator probes. For the glabrous skin, the tips of the toes were removed, and the skin was peeled back up to the ankle. The ankle was circumferentially cut, and the sciatic nerve was freed at hip level. The foot with the tibial nerve was placed in the organ bath chamber with superfused 30°C SIF buffer. Remaining muscle tissue, bones, and tendons were removed, and the skin was fixed in the "outside-out" configuration. The nerve was guided through a small opening to an adjacent chamber filled with mineral oil, where fine filaments were teased from the nerve and placed on a silver wire recording electrode.

Mechanically sensitive units were initially identified using blunt stimuli applied with a glass rod. Classification of mechanoreceptors was based on spike pattern and sensitivity to stimulus velocity, as described previously.[10–12] Raw electrophysiological data were recorded using a Powerlab 4/30 system and Labchart 8 software, with the spike-histogram extension supported by an oscilloscope for visual identification and subsequent analysis. All mechanical responses analyzed were adjusted for the latency delay between the electrical stimulus and the arrival of the action potential at the electrode.

Conduction velocity (CV) was measured using the formula CV = distance/time delay, where CVs > 10 m/s were classified as rapidly adapting mechanoreceptors (RAMs) or slowly adapting mechanoreceptors (SAMs), CVs < 10 m/s as Aδ fibers, and CVs < 1.5 ms/ as C-fibers.

Mechanically sensitive units were stimulated using a piezo actuator (Physik Instrumente (PI) GmbH & Co. KG, model P-841.60) connected to a force sensor with a calibrated conversion factor of Volts to Millinewtons. Various mechanical stimulation protocols were employed to identify and characterize sensory afferents. Vibrating stimuli with increasing amplitude and 20Hz frequency were applied to all types of mechanoreceptors, and the force required to evoke the first action potential was measured. A ramp and hold step with constant force (100 mN) was repeated with varying probe movement velocities (0.075, 0.15, 0.45, 1.5, and 15 mm s−1), and only firing activity during the dynamic phase was analyzed. SAM mechanoreceptors, Aδ mechanoreceptors, and nociceptors were mechanically tested with a constant ramp (1.5–2 mN m/s) and hold (10 seconds of static phase) stimulation, analyzing spikes evoked during the static phase. For electric search experimetns, data were collected from 5 *Tenm4*$^{fl/fl}$ (control) and 7 *Tenm4*$^{CKO}$ mice and from 4 WT (control) and 5 *Tenm4*$^{KI/+}$ mice. For the stimulus-response-function recordings, data were collected from 4 WT and *Tenm4*$^{KI/+}$ mice and 6 *Tenm4*$^{fl/fl}$ and 9 *Tenm4*$^{CKO}$ mice.

For the PreScission application, the five types of receptors are categorized into two main groups: mechanoreceptors (RA, SA, and D hairs) and nociceptors (AM and C fibers). Mechanoreceptors are subjected to a 20Hz step of increasing vibration amplitude applied at 30-second intervals, while nociceptors undergo a 150mM suprathreshold displacement at 30-second intervals. This protocol commences after isolating a receptive field with a metal ring. Prior to stimulation, it is crucial to verify the sealing status of the ring. The protocol involves stimulation for 3 minutes before applying the PreScission protease solution (concentration 20% (v/v) diluted in SIF). Stimulation is continued for an additional 45 minutes. At the end of the protocol, the conduction velocity is measured again to confirm the receptor's continued functionality. Experiments were conducted in a blinded manner (R.G. or M.A.K. were aware of the group allocations). To test for toxic (indirect) effects of cleavage products generated by the PreScission enzyme, single-unit recordings were performed on wild-type mechanoreceptors with receptive fields isolated using a metal ring. Simultaneously, skin from PreScission knock-in mice was treated with PreScission protease, and the resulting supernatant, containing cleavage products, was applied to the receptive fields of wild-type afferents. The responses of wild-type mechanoreceptors to repeated mechanical stimulation were monitored for up to 45 minutes.

**Electron microscopy**

Saphenous nerves were dissected from four 12-week-old *Tenm4*$^{CKO}$ and WT mice. The freshly isolated nerves were fixed in 4% paraformaldehyde/2.5% glutaraldehyde in 0.1 M phosphate buffer for 48 hours at 4°C. Following treatment with 1% OsO4 for 2 hours, each nerve underwent dehydration in a graded ethanol series and propylene oxide before being embedded in polyresin. Nerves were sectioned using a microtome into semi-thin (1µm) and ultrathin sections (70 nm). Semi-thin sections were stained with toluidine blue, and ultrathin sections were contrasted with uranyl acetate and lead acetate. Examination of semi-thin sections under a light microscope was performed to determine the total number of myelinated axons within the nerve. Ultrathin sections were examined using a Zeiss 910 electron microscope, and images were captured at an original magnification of 2500×. The A and C fiber dimensions were analyzed [6] using iTEM software.

**Behavior**

Von Frey filament testing was conducted to assess touch sensitivity in the mice. Animals were acclimated to the testing environment for two days in the week prior to the experiment. Paw withdrawal thresholds were then assessed using a standard Semmes-Weinstein set of von Frey filaments (ranging from 0.07 to 4 g, Aesthesio) in accordance with the up-and-down method.[13] The 50% paw withdrawal threshold was determined using the open source software, Up-Down Reader (https://bioapps.shinyapps.io/von_frey_app/).[14] For the ascending dose-response experiments, von Frey filament testing involved assessing each filament five times and calculating the responses as proportions. The cotton swab test measured sensitivity to gentle touch (0.7 – 1.6 mN), while brush testing evaluated dynamic mechanical sensitivity and measured intermediate touch sensation (>4 mN). A cotton swab (puffed out to approximately three times its original volume) and a soft brush (number 0) were applied to the plantar surface of the hind paw with gentle strokes. Each hind paw underwent testing three times, with a minimum interval of 5 minutes between consecutive tests. The number of withdrawals throughout the trials was counted and coded as a percentage of the total trials.

For Hargreaves assay, mice were acclimated for 30 minutes before the test. The infrared light source (Ugo Basile, 37370) was applied to the hind paw plantar surface, and withdrawal latency was automatically recorded. The focused radiant heat induced withdrawal within 10-15 seconds, with a 20-second cutoff to prevent tissue damage. The time required to withdraw the paws were recorded as indicators of their ability to assess temperature.

Sample sizes, described in each figure legend, were determined using GPower 3.1 software based on previous studies. No animals were excluded from the analysis. The mice were aged between 8 and 15 weeks old and were randomly assigned to various experimental groups, including control and other groups, ensuring a balanced representation across conditions. The experimenter was blinded to the group assignments during the von Frey filament testing.

Clasping behavior was assessed by suspending mice by the tail for 10 seconds and observing hindlimb posture. Normal behavior was defined as full hindlimb extension, while clasping was identified as hindlimbs retracting towards the body. Representative images were captured to illustrate hindlimb extension in control mice and hindlimb clasping in $Tenm4^{CKO}$ mice. Observations were scored, and the incidence of clasping behavior was analyzed for statistical significance.

The integrity of balance and coordination was assessed using the Rotar-rod system (83x91x61 - SD. Instruments, San Diego). This test is utilized to measure motor coordination and balance in mice. Mice were subjected to three trials, with 5-minute inter-trial intervals. Rotarod acceleration was set to 20 rpm over 240 seconds. The latency to fall (in seconds) was recorded, and the average of the three trials was calculated as an index of motor coordination and balance.

**Tripartite split-GFP complementation assay**

The tripartite split-GFP complementation assay[15] was performed in HEK293T cells (ACC 635, DSMZ) cultured in DMEM (DMEM 41966) supplemented with 10% FCS and 1% Penicillin/Streptomycin. When HEK293T cells reached approximately 70% confluence, they were co-transfected with GFP1-9::iRFP702 (Addgene #130125), mouseTenms(N-terminus)GFP10::mCHERRY, and mouseElkin1-GFP11::eBP2 plasmids using Fugene HD, following the manufacturer's instructions (Promega). Seven hours post-transfection, the culture medium was replaced with fresh medium. After 40 hours post-transfection, cells were fixed in 4% PFA, and images (eGFP, mCherry, and eBFP2) were acquired using cytationC10 (Biotek) with a 20x objective. Gen5 (Biotek) was used to analyze eGFP (interaction signal), mCherry (transfection signal of bait), and eBFP (transfection signal of prey) percentages of cells and fluorescence intensity. All signal intensities were normalized against background before color threshold overlapping for eGFP-positive (eGFP/mCherry) cell quantification.

**Cellular co-localization assay**

Transiently transfected N2a cells were imaged using an Airyscan fluorescence microscope (Axio, Zeiss) equipped with a ×63 1.4 NA oil lens. For cells transfected with either pRP TENM4-ALFA, pRK8 mScarlet-ELKIN1-EGFP, or mScarlet-PIEZO2, cells were prepared as previously described. After 48 hours of transfection, cells were fixed and stained with FluoTag®-X2 anti-ALFA, as described earlier. Cells were imaged in a total of four to five individual images under three experimental replicates. For each individual image, cells expressing both plasmids at similar expression levels were chosen and counted based on ALFA tag staining and mScarlet signals. The signal intensity in each group was normalized against control cells carrying only a single signal. The occurrence of normalized signals was statistically analyzed using Manders' Overlap Coefficient (MOC) as a colocalized indication built into ZEN.

**In vitro ALFA pull-down**

HEK293T cells were transfected with either Full-length C-terminally ALFA-tagged TENM4 coupled with ELKIN1 or ALFA plasmid coupled with ELKIN1. Membrane protein complexes were solubilized from total lysates prepared from RIPA buffer (50 mM Tris, pH 7.5, 150 mM NaCl, 1.0% NP-40, 0.5% deoxycholate, 0.1% SDS, 3 mM DTT) with 1% DDM for 1 hour on ice. Both lysates were incubated with 20µL of ALFA SelectorPE resin for 1 hour at 4 °C on a roller drum. After washing in PBS containing 0.3% DDM, bound proteins were eluted under native conditions by sequentially incubating twice with 50µL PBS containing 200µM ALFA peptide for 30 minutes at 4°C. Samples corresponding to 1/500 of the input and 1/50 of eluate fractions were resolved by SDS–PAGE. Analysis was performed by Western blotting using a polyclonal rabbit-raised antibody against ELKIN1 or RFP and a polyclonal sheep-raised antibody against TENM4 (dilution 1:500), followed by HRPconjugated goat anti-rabbit and anti-sheep IgG (Abcam Lot: GR288027-2, R&D Lot: 1522051, dilution 1:10,000). Blots were developed using the SuperSignal™ West Dura ECL Kit (Thermo Scientific) and imaged using a Bio-Rad ChemiDoc™ MP imaging system. Affinity purification of PIEZO2-mScarlet and TENM4-ALFA are performed as above.

**Microscopy setups**

Epifluorescence images were obtained with an LSM880 Airyscan microscope (Carl Zeiss GmbH), equipped with a 63× NA1.4 oil immersion lens, and operated the Airyscan detector in super-resolution mode. This results in a pixel size of 35 nm. Excitation laser lines are 488, 561, and 680 nm. Independent tracks are used to switch between lines with a Z-stack according to the sample thickness in 0.2 um spacemen. Each line of the image is first scanned at 488 nm or 561 nm and subsequently at 561 nm. Pixel dwell times are adjusted to a value between 1 and 2 microseconds with fourfold line averaging. After acquisition, Airyscan processing with default (=auto) parameters is employed, which results in a 16-bit image. This is stored in the proprietary CZI file format. data processing and signal quantification were performed using ImageJ.

**Mass spectrometry analysis**

The cell lines analysed were control Neuro2a cells or Neuro2a cells with a *Piezo1* genomic deletion as described previously [7]. The samples were lysed using 1% (w/v) sodium dodecyl sulphate and 1% (v/v) NP40, followed by SP3-bead-based digestion in an automated process utilising the AssayMAP Bravo robotic system (Agilent Technologies; adapted from).[16] In brief, protein lysates were treated with 5 mM dithiothreitol (DTT; Sigma-Aldrich) and heated to 90°C for 10 minutes for reduction. Alkylation was performed at room temperature with 10 mM iodoacetamide for 30 minutes, followed by quenching with 20 mM DTT for an additional 3 minutes. A mixture of 1 mg paramagnetic beads, comprising Sera-Mag Speed Beads, CAT# 09-981-121, and CAT# 09-981-123 (Thermo Fisher Scientific), was added, along with acetonitrile to a final concentration of 70% (v/v), and incubated for 18 minutes at room temperature. Beads were washed thrice with 200 µl 70% (v/v) ethanol, with 3-minute incubation at each step. Subsequently, 150 µl HEPES-KOH (pH 7.6) was added, followed by 4 µg of sequencing-grade LysC (Wako) and trypsin (Promega) for overnight incubation at 37°C. The supernatant was acidified with formic acid, and peptide solution was desalted using the AssayMap protocol. For reversed-phase liquid chromatography coupled to mass spectrometry (LC-MS) analysis, 1 µg of peptides per sample replicate was injected into an EASY-nLC 1200 system (Thermo Fisher Scientific) for separation employing a 110-minute gradient. Mass spectrometric measurements were conducted using an Exploris 480 (Thermo Fisher Scientific) instrument operating in data-independent acquisition (DIA) mode. Raw files were analysed using DIA-NN version 1.8.1,[17] in library-free mode, with an FDR cutoff 1195 of 0.01 and relaxed protein inference criteria whilst using the match-between runs option. Spectra were matched against a Uniprot mouse database (2022-03), including isoforms and a common contaminants database. Subsequent downstream analysis was performed using R version 4.3.2. MaxLFQ normalised protein intensities were log2 transformed and filtered to retain at least 4 valid values in at least one group for each protein before applying a downshift-imputation procedure. Significance determination utilised the limma package,[18] to compute two-sample moderated t statistics, with nominal P-values adjusted using the Benjamini-Hochberg method.

Quantitative analysis of distinct peptides was conducted by utilising the report.tsv output generated by DIA-NN, followed by application of filtering criteria wherein peptides with Protein.Q.Value and Lib.Q.Value both less than or equal to 0.01 were retained for further analysis. Subsequently, the sum of the three most abundant fragments was computed for each precursor. The precursor with the highest abundance (Top1) was then selected for each distinct peptide sequence to facilitate peptide-level quantitation. Peptide maps were constructed by calculating position-based intensities, obtained by dividing the intensity of each peptide sequence by its respective length. In cases where multiple sequences of the same stretch were identified, intensities were aggregated to account for miscleavages or overlapping peptide sequences. Peptide intensities are not normalised. The mass spectrometry proteomics data have been deposited to the ProteomeXchange Consortium via the PRIDE partner repository with the dataset identifier PXD050952.[19]

**Immunogold labeling and transmission electron microscopy**

Dissociated DRG neurons were fixed by adding an equal volume of 8% Paraformaldehyde Aqueous Solution (PFA, pH 7.4, pre-warmed to 37 °C) in 2x PBS to the culture medium to obtain a final concentration of 4% PFA and 1x PBS. After 15 min at room temperature, the fixative was replaced with fresh 4% PFA in 1x PBS, and samples were incubated for an additional hour at 4°C. Following three washes in PBS, neurons were blocked in 10% normal goat serum in PBS 1x for 30 min at room temperature and incubated overnight at 4 °C with the TENM4-C-terminal antibody diluted 1:200 or 1:500 in blocking solution. After three PBS washes, cells were incubated for 1h at room temperature with 12 nm or 6 nm Colloidal Gold AffiniPure® Goat Anti-Rabbit IgG (H+L) (1:30 dilution in blocking solution), followed by three final PBS washes. Labeled samples were then prepared for subsequent resin embedding and ultrathin sectioning for electron microscopy.

Following pre-embedding immunogold labelling, DRG neurons were fixed additionally in 4% PFA (v/v) and 2.5% glutaraldehyde (v/v) in 0.05 M HEPES for 2 hours at room temperature. Samples were processed using a modified version of the rOTO protocol by[20]. Samples were contrasted with 2% (v/v) Osmium Tetroxide Aqueous Solution and 1.5% (w/v) potassium ferrocyanide in 0.05 M HEPES for 60 minutes on ice, washed with 0.1% (w/v) thiocarbohydrazide for 10 minutes, followed by an incubation in 1% (w/v) osmium tetroxide for 30 minutes at RT. Final contrast was achieved by an incubation in 1% (w/v) uranyl acetate for 30 minutes at RT. After dehydration through a graded series of ethanol, embedding was done in Poly/Bed® 812 Embedding Media. Ultrathin resin sections (150 nm) of cells attached to the Lumox® membrane were collected on slot grids and stained with 3% lead citrate. Imaging was performed at 200 kV using the JEM 2100 Plus. Acquisition was done with the Xarosa camera and the Radius EM imaging software package.

Images were acquired for the experimental condition (TENM4-C-terminal antibody) and the negative control (no primary antibody). For the experimental condition, the total analyzed contact area was 176 $\mu m^2$, calculated as the measured cumulative neurite-laminin contact length multiplied by the section thickness (0.15 µm); a comparable contact area (>180 $\mu m^2$) was imaged for the negative control. Gold particles located within 200 nm of the neurite membrane were counted and classified as being at the neurite-laminin interface or elsewhere. Gold-labeled tethers were defined as electron-dense structures at the interface that were associated with at least one gold particle. Tether density was calculated either relative to the total contact area or to the contact area of neurites with at least one gold particle. Neurites were classified according to the number of associated tethers, and tethers were further categorized based on the number of associated gold particles, yielding the distributions of tethers per neurite and gold particles per tether, respectively.

**Statistical analysis**

Statistical analyses were performed using GraphPad Prism (version X). The study employed a variety of tests for different experimental assessments. Chi-squared and Fisher's exact tests evaluated proportions of neuron types, Mann-Whitney tests compared various responses, and Two-way repeated measures ANOVA with Sidak's post-test assessed stimuli-induced firing activity. Additionally, the impact of PreScission protease treatment was scrutinized with Friedman tests, Dunn's multiple comparisons tests, and Fisher's exact tests. The strength of interaction between ELKIN1 and TENM4 was determined via One-way ANOVA. A comprehensive statistical strategy, including Kruskal-Wallis tests, was implemented to analyze various datasets. Steps were taken to ensure that control and experimental groups were spread across different days to reduce bias.

Figure S1

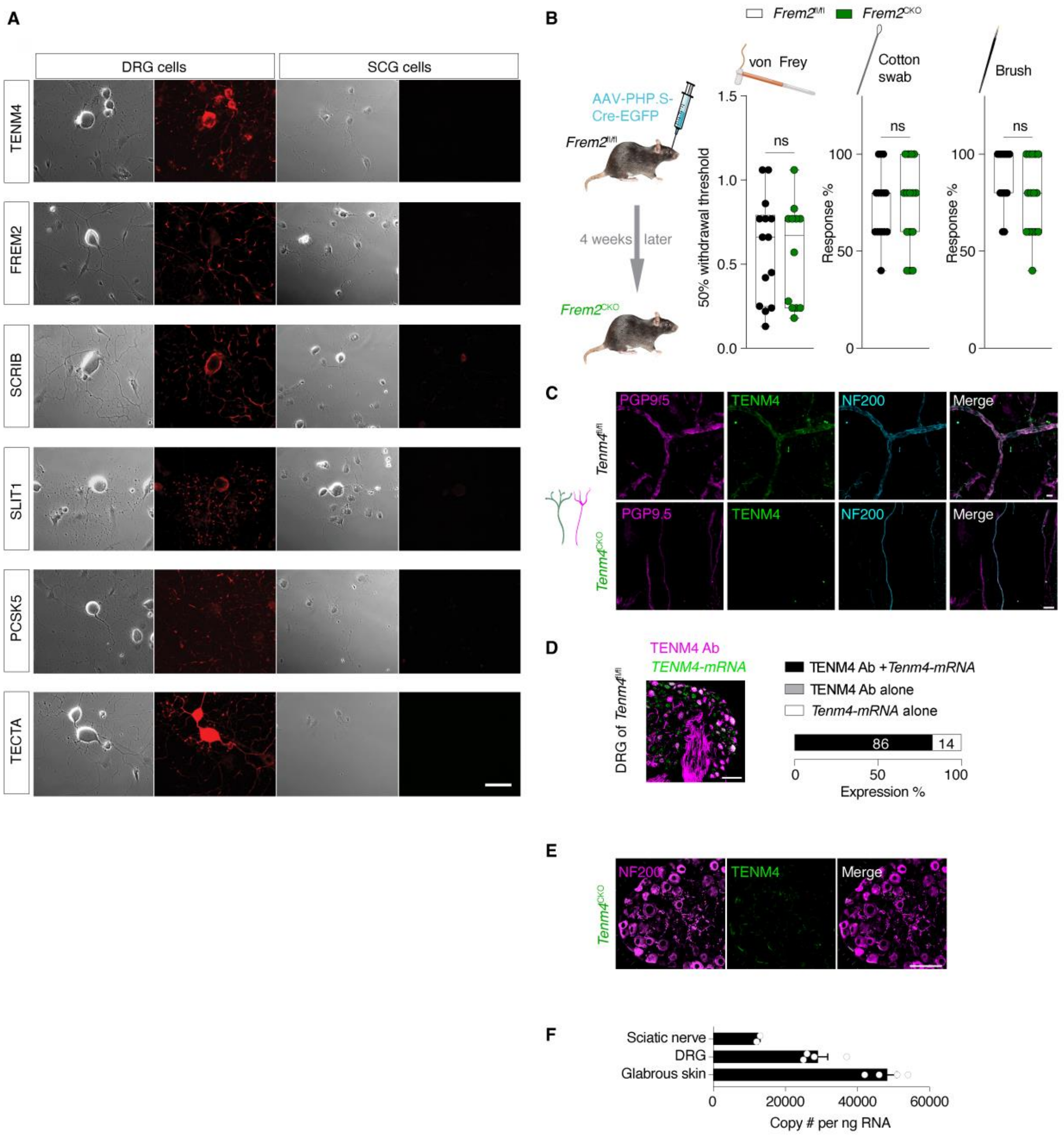


**Figure S1. Tether candidate screen and validation of TENM4 deletion.** (**A**) The 6 tether protein candidates exhibit a patchy-like expression pattern in the neurites of DRG cells, and no staining was observed in the non-mechanosensitive SCG cells cultured on plates coated with laminin. Scale bar is 10μm. (**B**) Left, a diagram depicting generation of *Frem2*$^{CKO}$ mice via retro-orbital injection of *pAAV-Syn-Cre-p2A-EGFP* in the sinus of *Frem2*$^{fl/fl}$ mice. Right, *Frem2*$^{CKO}$ mice exhibit normal touch sensitivity compared to the control *Frem2*$^{fl/fl}$ mice. (**C**) TENM4 positive staining found in nerve endings (in hairy skin), visualized by PGP9.5 (a neuronal marker) and NF200 in WT mice (top panels). *Tenm4*$^{CKO}$ mice exhibit a notable decrease in TENM4+ sensory nerve endings (below panels). Scale bar 20μm. (**D**) Left, representative image demonstrates the colocalization of a smFISH probe targeting Tenm4 mRNA with the

TENM4 antibody staining. Right, quantification of TENM4+ sensory neurons stained with both the smFISH probe and the antibody. Scale bar is 100µm. Quantification in each group from 2 male and 3 female mice; more than 700 neurons were counted in each group. (**E**) Representative DRG sections showing absence of TENM4+ DRG cells in sections from *Tenm4*$^{CKO}$ mice, NF200 staining was unchanged, scale bar 100µm. (**F**) Quantification of *Tenm4* mRNA expression level (copies of transcript per ng total RNA) in skin, DRG, and sciatic nerve. Quantification in each group from 2 male and 2 female mice.

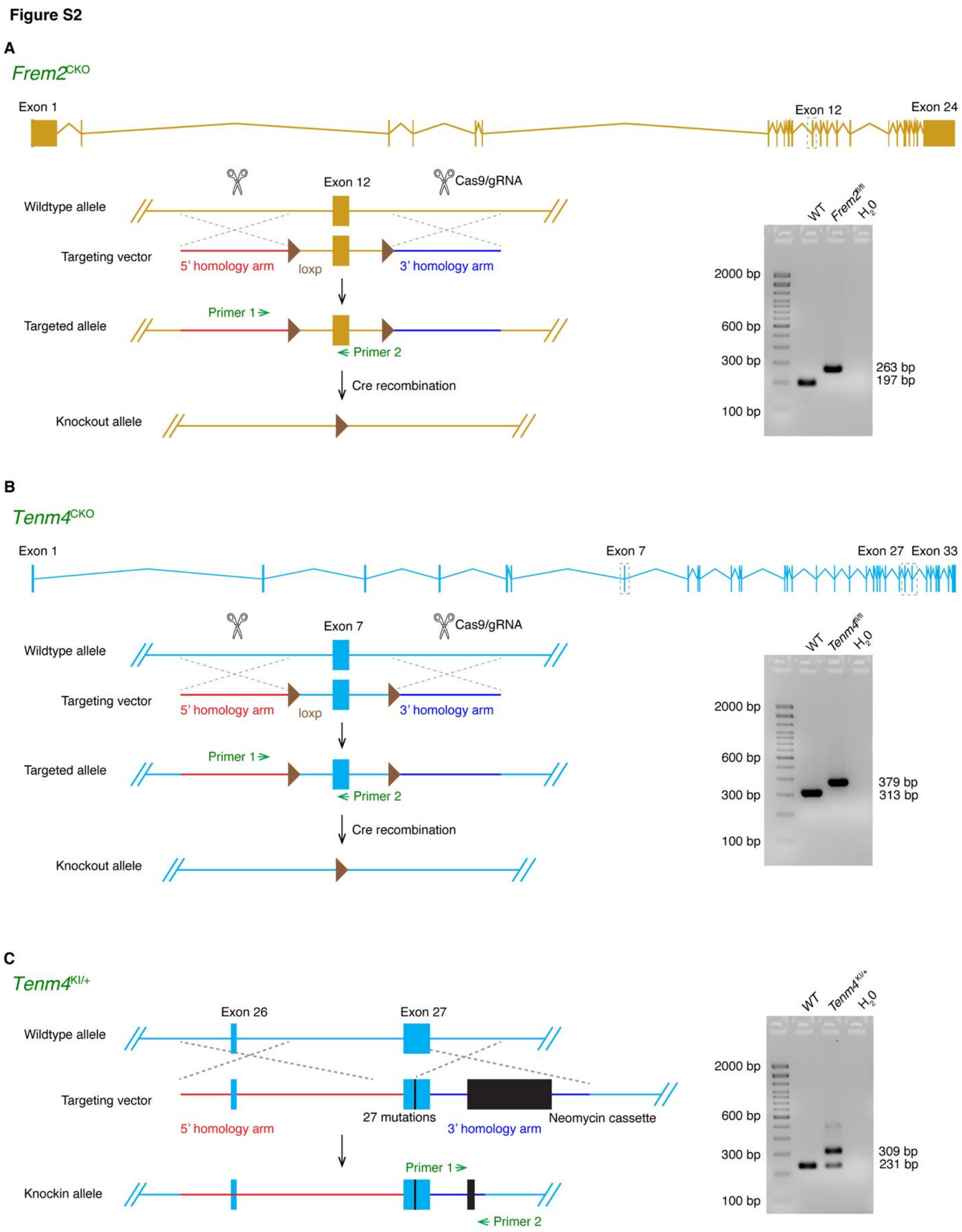


**Figure S2. Strategies of generating transgenic mice**. (**A**) Schematic of the *Frem2*$^{CKO}$ mouse model targeting strategy using a targeting vector constructed with Infusion technology. The plasmid structure includes a 5' homologous arm (3.0kb), a LoxP sequence (1.0kb), and a 3' homologous arm (3.0kb). Right,

PCR bands from genomic DNA of WT, *Frem2*$^{fl/fl}$ are presented. (**B**) Diagram depicting *Tenm4*$^{CKO}$ mouse model creation with an Infusion-constructed targeting vector (5' homologous arm: 3.0 kb, flox: 0.9 kb, 3' homologous arm: 3.0 kb), along with PCR bands from genomic DNA of WT and *Tenm4*$^{fl/fl}$ mice. (**C**) Diagram illustrating the creation of the *Tenm4*$^{KI/+}$ mouse model using homologous recombination in ES cells. The targeting vector was designed to introduce a 21 bp change in exon 27 of Tenm4 (5' homologous arm: 7.9 kb, and 3' homologous arm: 2.5 kb) and the neomycin selection cassette, flanked by FRT sites. The neomycin cassette excision was performed by crossing the resulting chimeric mice with C57BL/6 FLP mice, leaving one FRT site (78 bp), which was used for genotyping Tenm4 KI mice. On the right, PCR bands from genomic DNA of WT and *Tenm4*$^{KI/+}$ mice are displayed.

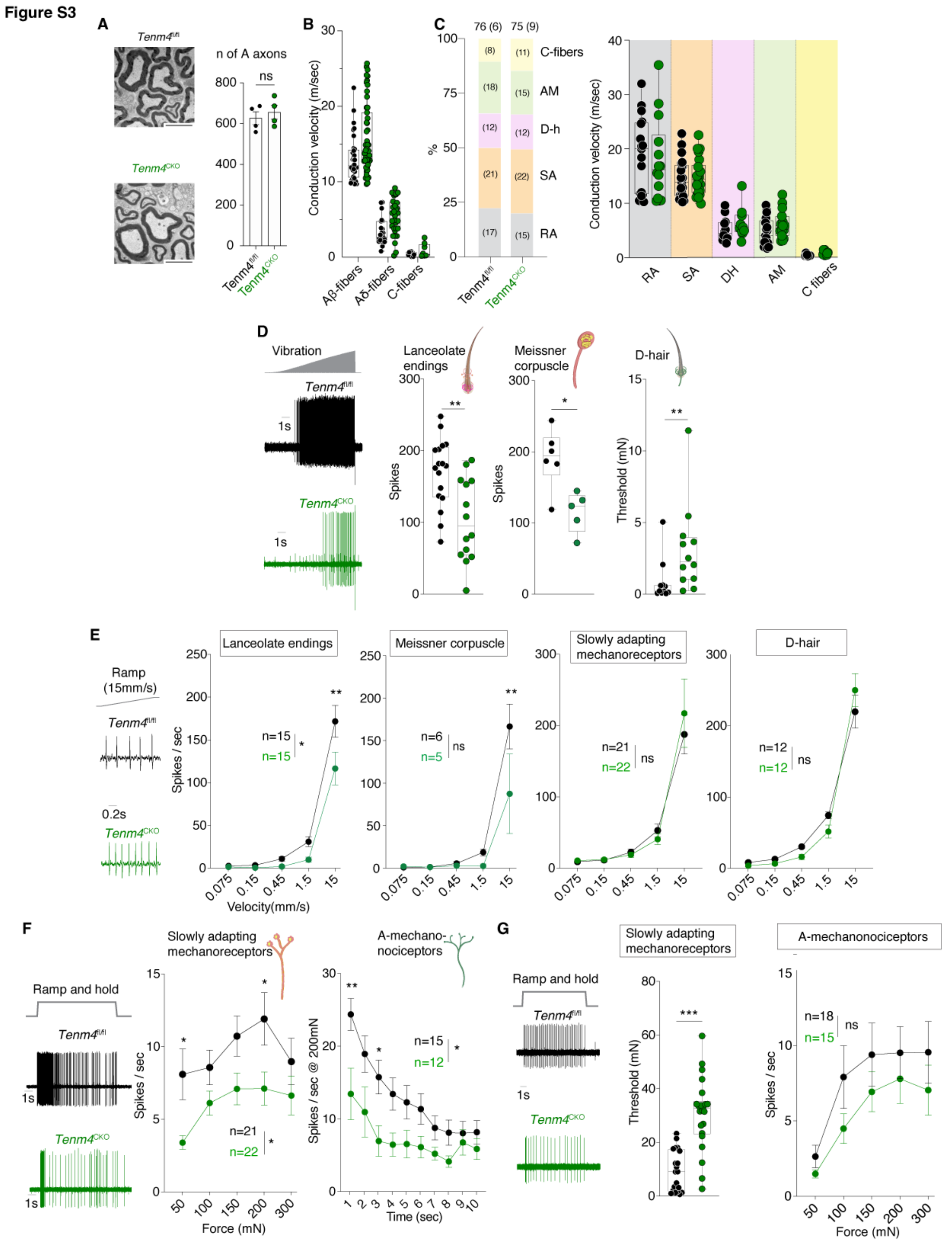


**Figure S3. Stimulus-response functions after *Tenm4* gene deletion.** (**A**) Example electron-micrographs from the saphenous nerve of a *Tenm4*$^{fl/fl}$ (top) and *Tenm4*$^{CKO}$ mouse (bottom) with quantification of axon number in each case. Scale bar is 20µm. (**B**) Conduction velocities of sensory afferents recorded from 6 *Tenm4*$^{fl/fl}$ and 9 *Tenm4*$^{CKO}$ mice. (**C**) Proportions and conduction velocities of different sensory afferents recorded from 6 *Tenm4*$^{fl/fl}$ and 9 *Tenm4*$^{CKO}$ mice. (**D**) Example traces of vibration sensitive RAMs. Firing to vibration stimuli in RAMs associated with hairs or Meissner's corpuscles in *Tenm4*$^{CKO}$ mice were significantly reduced. RAM Aδ-fibers innervating D-hairs in *Tenm4*$^{CKO}$ mice exhibited significantly higher mechanical thresholds compared to control mice. Significance calculated with Mann-Whitney test. **(E**) Example traces of single sensory units from *Tenm4*$^{fl/fl}$ and *Tenm4*$^{CKO}$ mice responding to a ramp stimulus with a 15 mm/sec movement velocity. The firing activity of sensory afferents innervating Merkel cells and D-hairs, in response to varying movement velocities, showed no significant differences between *Tenm4*$^{fl/fl}$ and *Tenm4*$^{CKO}$ mice. Two- way repeated measures ANOVA followed by Sidak's post-test; ns indicates $p >$ 0.05. Data represent the mean ± s.e.m and were collected from 6 *Tenm4*$^{fl/fl}$ and 9 *Tenm4*$^{CKO}$ mice. (**F**) Example SAM responses to a 50 mN amplitude ramp and hold stimulus. Stimulus response functions of SAMs were significantly impaired in *Tenm4*$^{CKO}$ mice. Additionally, Aδ-mechanonociceptors showed reduced firing during a 10s long 200 mN stimulus compared to control mice. Significance calculated with two-way repeated measures ANOVA followed by Sidak's post-hoc tests. Data collected from 6 *Tenm4*$^{fl/fl}$ and 9 *Tenm4*$^{CKO}$ mice. (**G**) Example traces of single sensory afferents innervating Aδ mechanonociceptors in in *Tenm4*$^{fl/fl}$ and *Tenm4*$^{CKO}$ mice responding to a ramp-and-hold stimulus with a constant force of 100mN. The mechanical threshold and firing activity of various sensory afferents showed no significant difference between *Tenm4*$^{fl/fl}$ and *Tenm4*$^{CKO}$ mice. Mann-Whitney test and Two-way repeated measures ANOVA followed by Sidak's post-test; ns indicates $p > 0.05$. The line graph represents the mean ± s.e.m. and were collected from 6 *Tenm4*$^{fl/fl}$ and 9 *Tenm4*$^{CKO}$ mice.

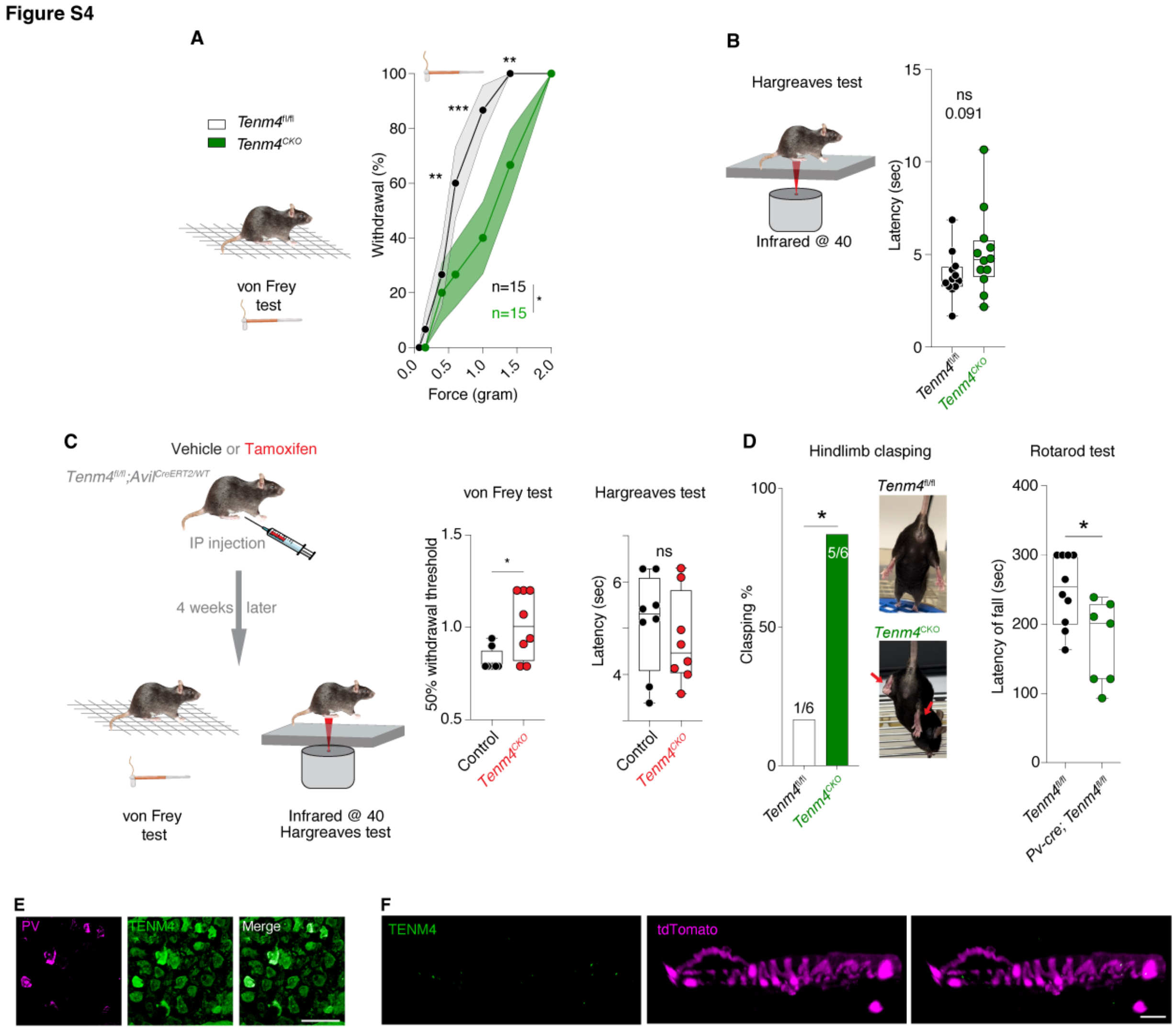


**Figure S4. *Tenm4*$^{CKO}$ mice display mechanosensory and proprioceptive deficits but normal responses noxious heat.** (**A**) Responses to an ascending series of von Frey hair stimuli in *Tenm4*$^{CKO}$ mice. The line graph illustrates the mean represented by points and the surrounding shades indicate s.e.m. (**B**) Left, a Hargreaves test schematic for paw withdrawal latency, indicating thermal pain thresholds. Right, *Tenm4*$^{CKO}$ mice showed comparable responses to heat stimulus, similar to *Tenm4*$^{fl/fl}$ mice. (**C**) Left, a diagram illustrating the generation of *Tenm4*$^{CKO}$ and control mice via tamoxifen injections in *Tenm4l*$^{fl/fl}$;*Avil*$^{CreERT2/WT}$ mice. Right, *Tenm4*$^{CKO}$ mice exhibited significantly higher withdrawal thresholds to von Frey stimuli compared to control mice (left), while comparable responses to heat stimulus, similar to the control mice (right). Mann-Whitney test; ns indicates $p > 0.05$. Data were collected from 12 (a), 8 (b), animals per group. Means ± s.e.m. (**D**) Left, A significant increase in clasping behavior is observed in *Tenm4*$^{CKO}$ mice, compared to controls (*Tenm4*$^{fl/fl}$). Representative images show normal hindlimb extension in *Tenm4*$^{fl/fl}$ mice (top) and hindlimb clasping in *Tenm4*$^{CKO}$ mice (bottom, indicated by red arrows). Right, *Pv*$^{Cre}$; *Tenm4*$^{fl/fl}$ mice show a significant reduction in latency to fall, indicating impaired motor coordination. Each point represents an individual mouse. (**E**) Representative DRG sections showing double labeling of TENM4 and PV. Scale bar: 100 µm. All PV$^{+}$ cells were also TENM4$^{+}$. Two male and three female mice were used. (**F**) TENM4 expression in tdTomato-labeled muscle spindle and Golgi tendon organ afferent terminals in muscle of *PV*$^{Cre}$;*Rx3*$^{Flpo}$;Ai65D mice.

**Figure S5**

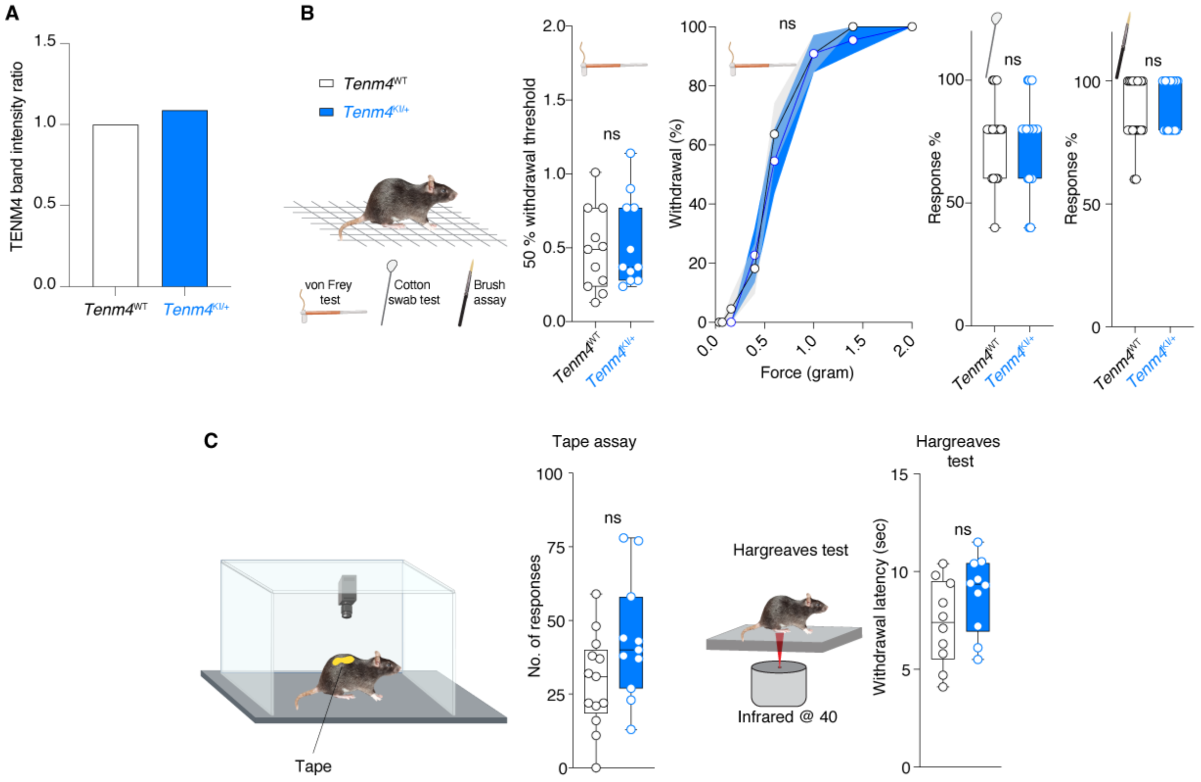


**Figure S5. Behavioral characterization of *Tenm4*$^{KI/+}$ mice.** (**A**) *Tenm4*$^{KI/+}$ mice exhibit comparable TENM4 expression to wildtype mice, indicating that the knock-in mutation does not alter overall protein abundance. (**B**) *Tenm4*$^{KI/+}$ mice exhibited similar withdrawal thresholds and responses to von Frey filaments, a cotton swab, and dynamic brush, compared to wildtype mice. (**C**) *Tenm4*$^{KI/+}$ mice display normal responses to tape and heat stimuli, similar to WT mice. Mann-Whitney test and Two-way repeated measures ANOVA followed by Sidak's post-test; ns indicates $p > 0.05$. Data were collected from 11 (von Frey, cotton swab and brush tests), 13 (tape assay) and 10 (Hargreaves test) mice in each category. The line graph represents the mean ± s.e.m.

Figure S6

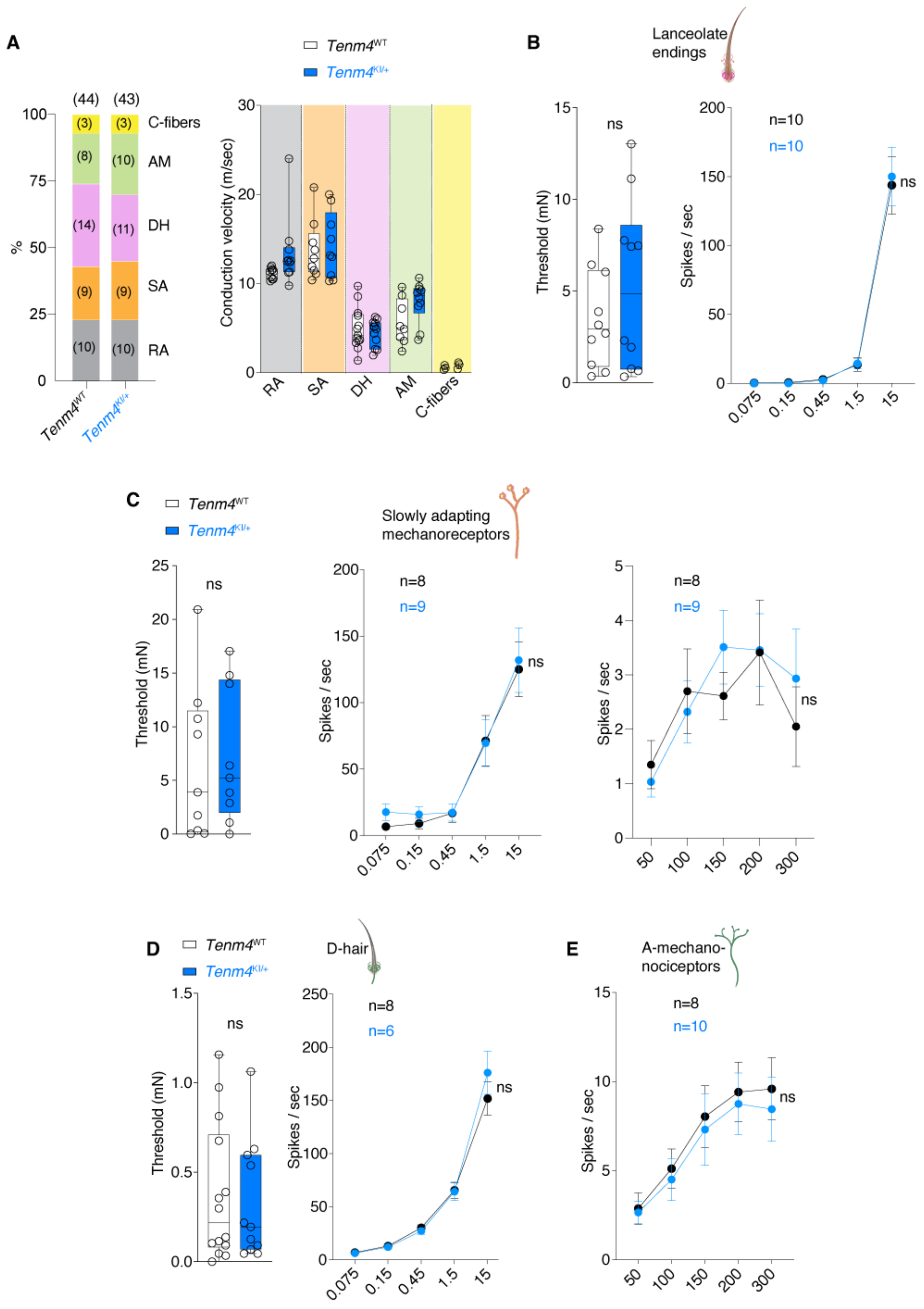


**Figure S6. Electrophysiological characterization of *Tenm4*$^{KI/+}$ mice.** (**A**) Proportions and conduction velocities of different sensory afferents recorded from 5 WT and 4 *Tenm4*$^{KI/+}$ mice show no marked changes. (**B**) The mechanical threshold and firing activity during the dynamic phase of ramp stimuli showed no significant differences between RAMs in WT and *Tenm4*$^{KI/+}$ mice. (**C**) The mechanical threshold firing activity during the dynamic phase of ramp stimuli, as well as the static phase of ramp-and-hold stimuli, showed no significant differences between SAMs in WT and *Tenm4*$^{KI/+}$ mice. (**D**) The mechanical threshold and the firing activity during the dynamic phase of ramp stimuli showed no significant differences between D hairs receptors in WT and *Tenm4*$^{KI/+}$ mice. (**E**) The firing activity during the static phase of ramp-and-hold stimuli, showed no significant differences between Aδ mechanonociceptors in WT and *Tenm4*$^{KI/+}$ mice. Mann-Whitney test and Two-way repeated measures ANOVA followed by Sidak's post-test; ns indicates $p > 0.05$. Data were collected from 5 WT and 4 *Tenm4*$^{KI/+}$ mice. Means ± s.e.m.

**Figure S7**

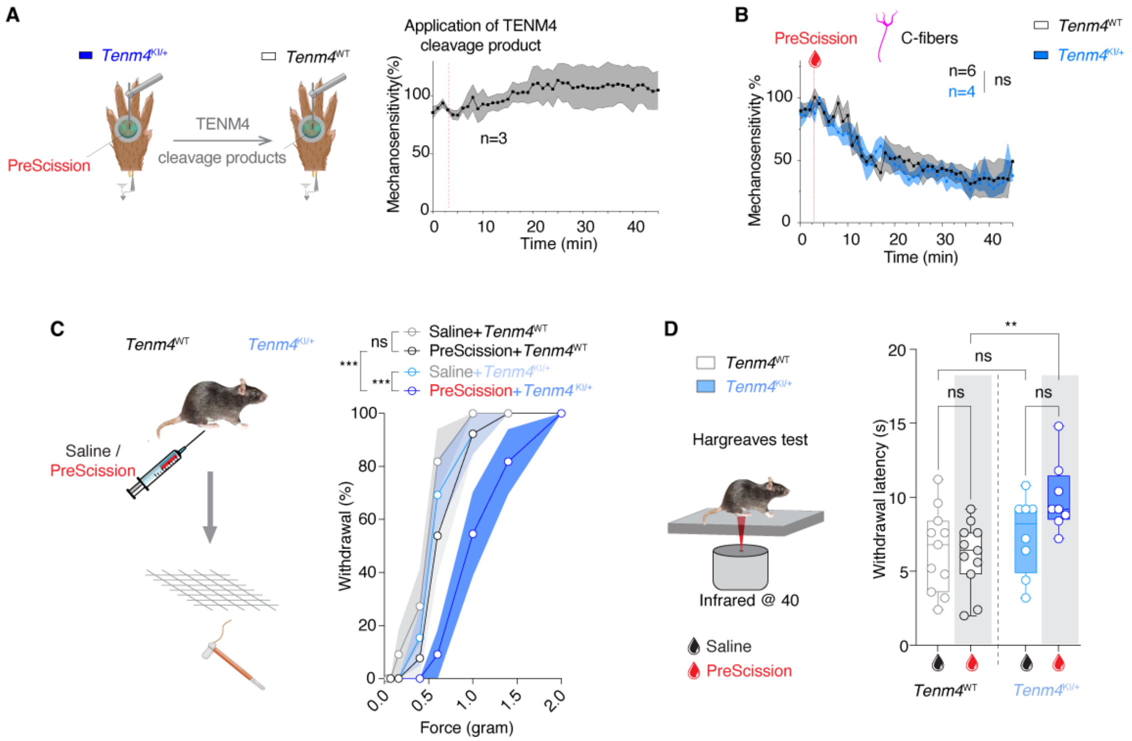


**Figure S7. TENM4 cleavage impairs touch sensitivity but not C-fiber function.** (**A**) Left, schematic showing fluid containing the cleavage product from PreScission-treated *Tenm4*$^{KI/+}$skin applied to the isolated receptive fields of WT mechanoreceptors. Right, mean mechanoreceptor responses (% of baseline) for RAM-Aβ fiber type in hairy skin, before and after the application of TENM4 cleavage product (n=3 units). Note that the mechanoreceptors retained mechanosensitivity during the 45 mins of observation. (**B**) C-fibers in WT and *Tenm4*$^{KI/+}$ mice retain functionality after exposure to PreScission protease applied to their receptive fields. (**C**) Behavioral assays used to assess the effects of TENM4 cleavage 1 hr after PreScission injection to the plantar hind paw. The line graph illustrates the mean represented by points and the surrounding shades indicate s.e.m. A dramatic decrease in sensitivity to von Frey stimulation was observed only after treatment of *Tenm4*$^{KI/+}$mice with PreScission protease, and not with saline. (**D**) Left, Schematic of the Hargreaves test Right, *Tenm4*$^{KI/+}$ mice injected with PreScission showed slightly longer withdrawal latencies (i.e., decreased responses) to heat stimulus, compared to PreScission-injected WT mice. Data were collected from 11 WT and 8 *Tenm4*$^{KI/+}$ mice. Two-way repeated measures ANOVA followed by Sidak's post-test; ns indicates $p > 0.05$; ** indicates $p < 0.01$.

**Figure S8**

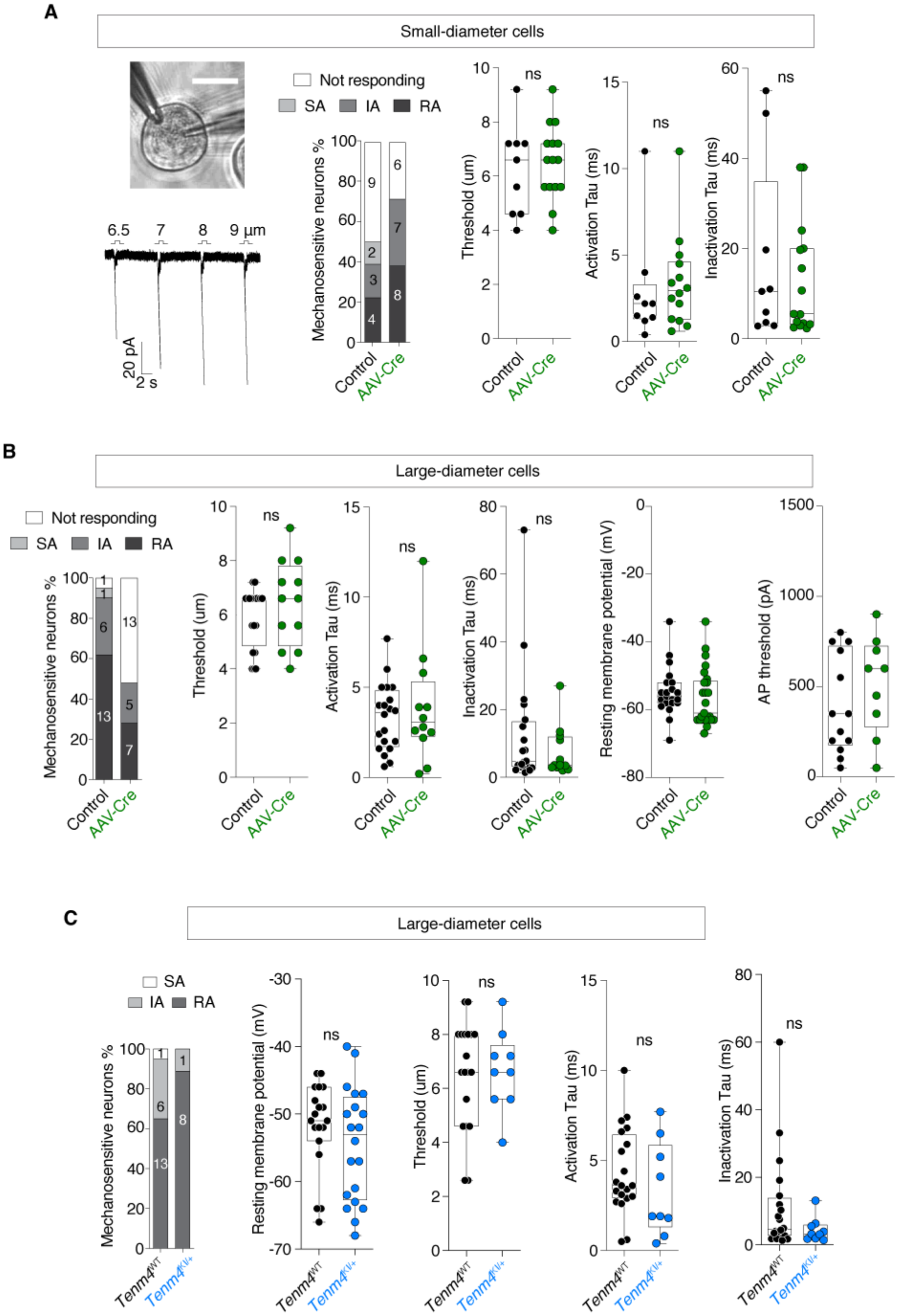


**Figure S8. Mechanically activated current kinetics of DRG neurons from *Tenm4*$^{CKO}$ and *Tenm4*$^{KI/+}$ mice.** (**A**) Representative traces of poking-induced currents resulting from indentations ranging between 1 and 9µm on GFP+ large-diameter neurons in *Tenm4*$^{CKO}$ mice. Small-diameter neurons from both *Tenm4*$^{fl/fl}$ and *Tenm4*$^{CKO}$ mice exhibited similar number of cells displaying rapidly, intermediate, and slowly adapting MA-currents, mechanical thresholds, as well as activation and inactivation kinetics. Chi- squared test and Mann-Whitney test; ns indicates $p > 0.05$. Data were collected from 5 *Tenm4*$^{fl/fl}$ and 5 *Tenm4*$^{CKO}$ mice. Means ± s.e.m. (**B**) *Tenm4*$^{CKO}$ mice had a deficit in MA-currents in large neurons compared to control *Tenm4*$^{fl/fl}$ mice. Large-diameter neurons from both *Tenm4*$^{fl/fl}$ and *Tenm4*$^{CKO}$ mice exhibited similar number of cells displaying rapidly, intermediate, and slowly adapting MA-currents, mechanical thresholds, activation and inactivation kinetics, as well as Resting membrane potential and action potential threshold. (**C**) Recordings from large diameter sensory in culture (scale bar = 20μm) showed a substantial decrease in the proportion of cells with poking-induced currents, specifically in cells from *Tenm4*$^{KI/+}$ treated with the PreScission protease. Significance calculated with Chi-squared test. Large-diameter DRG neurons from PreScission- treated WT and PreScission-treated *Tenm4*$^{KI/+}$ mice exhibited similar percentages of rapidly, intermediate, and slowly adapting MA-currents, mechanical thresholds, as well as activation and inactivation kinetics.

Figure S9

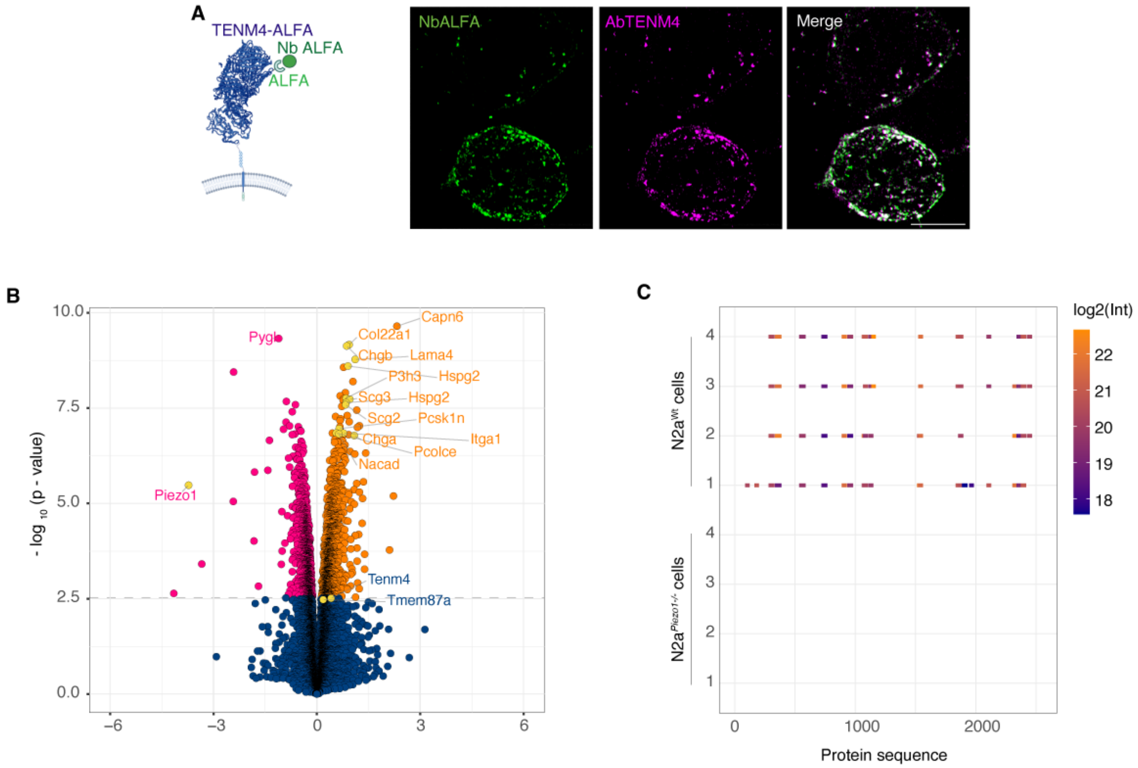


**Figure S9. Generation of ALFA tagged TENM4 and the proteomic profile of N2a cells and N2a*Piezo1-/-* cells**. (**A**) Left, schematic of ALFA tagged TENM4. The ALFA epitope should be recognized by anti-ALFA nanobody (NbALFA) without the need to permeabilize the cells. Right, N2a cells transfected with TENM4-ALFA and stained with NbALFA labeled with AbberiorStar635P (NbALFA-Ab635P, green) and our anti-TENM4 C-terminal antibody (purple). Note the clear colocalization of the two signals (white in merge). (**B**) Volcano plot of mass spectrometry analyses of N2a*Piezo1-/-* compared to WT N2a cells (n = 4). Both Elkin1 (Tmem87a) and TENM4 were endogenously expressed in N2a cells and both were slightly upregulated upon Piezo1 gene deletion. Many of the proteins extensively upregulated in N2a*Piezo1-/-* cells (highlighted in the plot) are involved either in the organization of the extracellular matrix (ECM) or in the regulation of the secretory granules, suggesting a link between mechanosensitivity and ECM composition. (**C**) Peptide map shows local peptide coverage of PIEZO1 in WT N2a cells. No PIEZO1 peptides were detected in the N2a*Piezo1-/-* samples. Displayed values are in linear space whilst a log2 transformation is applied to the color scheme.

Figure S10

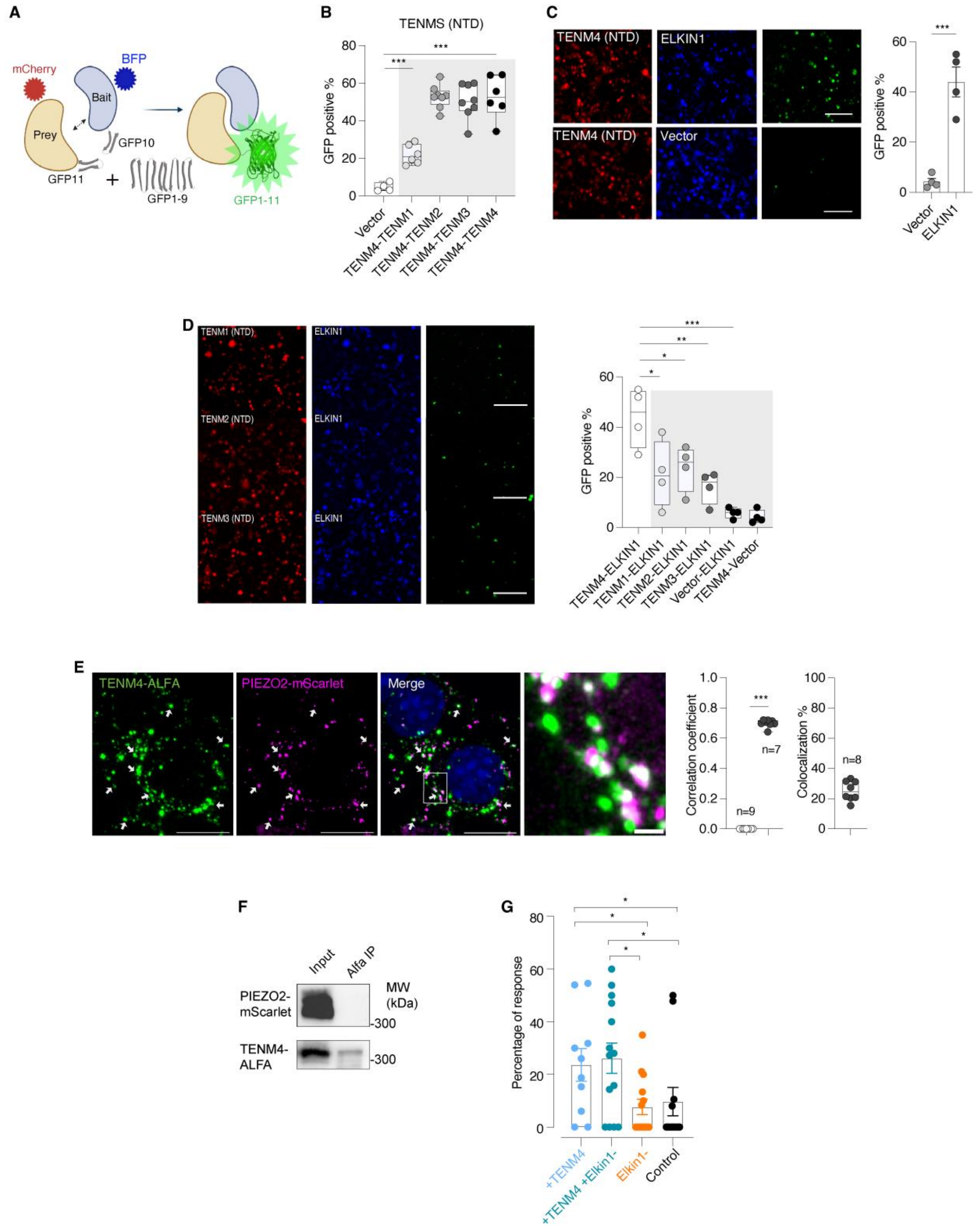


**Figure S10. TENM4 interacts with other Teneurins and ELKIN1.** (**A**) Schematic of the tripartite split-GFP complementation assay. (**B**) Quantification of the GFP-complementation signal in the split-GFP assay. Assay confirms heterologous interactions between the Teneurins. Pair wise interactions tested with

TENM4(NTD) between TENM1(NTD), TENM2(NTD), TENM3(NTD) and TENM4(NTD). (**C**) HEK293T cells were transfected with three plasmids encoding bait (e.g., ELKIN1)::GFP10, prey (e.g., TENM4)::GFP11 and GFP domain 1-9 proteins. Protein interaction produces complementation and reconstituted GFP fluorescence (green). Representative images showing complementation signals in cells expressing the N-terminal domain (NTD) of TENM4 showing a specific interaction with ELKIN1, not seen in the empty vector control (left 6 panels). Quantification (right) shows a strong interaction between TENM4(NTD) and ELKIN1. Scale bar 100 μm. Statistical significance calculated with Mann-Whitney test. (**D**) Representative images depicting the tripartite-GFP interaction between ElKIN1 and other Teneurins N-terminals (TENM1 (NTD), TENM2 (NTD), or TENM3 (NTD)). Scale bar is 100μm. Quantification (right) shows the interaction between ElKIN1 and other Teneurins. Statistical analysis with one-way ANOVA followed by Dunnett's multiple comparison test. * indicates $p < 0.05$; ** indicates $p < 0.01$; *** indicates $p < 0.001$ means ± s.e.m. (**E**) Representative super-resolution images of N2a cells co-transfected with a *Tenm4-ALFA* expression plasmid with PIEZO2-mScarlet (purple), arrows indicate co-localization signals. Right higher magnification of co-localized protein region (from white dash boxes) Scale bar is 10μm for cells, 1μm magnified region. Quantification was carried out and a co-localization correlation, ranging from -1 to 1, signifies the strength of overlap. Values near 1 indicate strong co-localization, near -1 indicate separation. Right, TENM4 shows 30% with PIEZO2 on the cell surface. (**F**) HEK293T cells were co-transfected with PIEZO2-mScarlet and TENM4-ALFA. Cell lysates (Input) and proteins eluted from ALFA selector resin (Alfa IP) were analyzed by Western blot using antibodies against mScarlet and ALFA. PIEZO2-mScarlet was not detected in the ALFA pull-down together with TENM4-ALFA, indicating the interaction might not be direct. Molecular weight markers are indicated (kDa). (**G**) Percentage of cells exhibiting mechanically evoked responses to pillar deflection under the indicated conditions. N2a cells were transfected with Tenm4-ALFA, Elkin1 siRNA, both, or control siRNA. Dots represent individual cells from 3 mice; bars show mean ± SEM. $p < 0.05$.

Figure S11

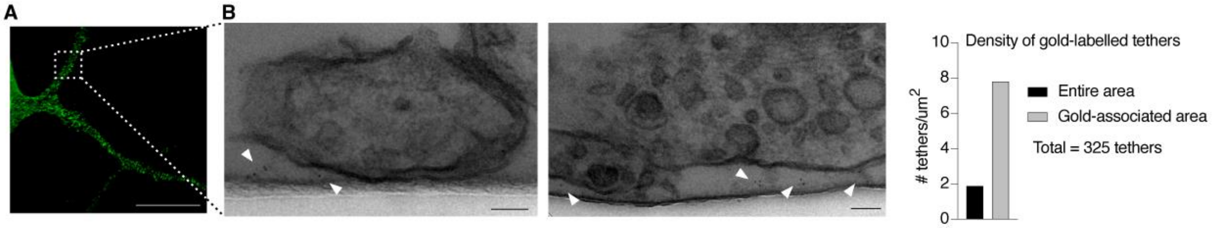


**Figure S11. TENM4 is located at sensory tethers.** (**A**) Immunofluorescence image of a mouse sensory neuron neurite stained with the Anti-TENM4-C antibody recognizing extracellular TENM4 epitopes, showing a punctate membrane distribution of TENM4 along the neurite. Scale bar: 20 µm. (**B**) Left, Zoomed-out images for the TEM images in Figure 7B provide overall neurite context. Scale bars: 100 nm. Right, density of gold-labeled tethers calculated using either the total neurite-laminin contact area or only the area of neurites associated with at least one gold particle.

**Table S1**

| Gene | DRG | | SCG | |
|---|---|---|---|---|
| | Mean | SEM | Mean | SEM |
| Adamts12 | N.E.D | | N.E.D | |
| Adamts20 | 0.163 | 0.069 | 0.068 | 0.037 |
| Apc2 | 0.177 | 0.013 | 0.037 | 0.015 |
| C3 | 0.232 | 0.021 | 0.3 | 0.018 |
| Celsr1 | 0.464 | 0.071 | 0.401 | 0.04 |
| Clstn2 | 0.673 | 0.034 | 0.276 | 0.08 |
| Coagulaion factor8 | 0.387 | 0.012 | 0.09 | 0.041 |
| Col5a1 | N.E.D | | 0.469 | 0.158 |
| Dmbt1 | N.E.D | | N.E.D | |
| Fbn2 | 0.322 | 0.085 | 0.308 | 0.038 |
| Frem2 | 0.362 | 0.062 | N.E.D | |
| Igsf10 | 0.482 | 0.044 | 0.336 | 0.006 |
| Madd | 0.405 | 0.151 | 0.391 | 0.119 |
| Notch3 | 0.602 | 0.072 | 0.309 | 0.023 |
| Pcnxl3 | 0.595 | 0.217 | 0.439 | 0.022 |
| Pcsk5 | 0.388 | 0.325 | N.E.D | |
| Pkhd1l1 | 0.361 | 0.008 | 0.109 | 0.016 |
| Plxnb1 | 0.66 | 0.053 | 0.541 | 0.002 |
| RIKEN 672066o15 | 0.823 | 0.137 | 0.34 | 0.049 |
| 4931403E03Rik | 0.103 | 0.016 | 0.178 | 0.028 |
| 5430411K18Rik | 0.497 | 0.071 | 0.359 | 0.041 |
| Robo1 | 0.414 | 0.069 | 0.378 | 0.102 |
| Scrib | 0.188 | 0.066 | N.E.D | |
| Slit1 | 0.234 | 0.025 | N.E.D | |
| Slit2 | 0.568 | 0.043 | 0.23 | 0.07 |
| Stab1 | 0.519 | 0.034 | 0.482 | 0.019 |
| Strc3 | N.E.D | | N.E.D | |
| Svep1 | 0.572 | 0.01 | 0.36 | 0.178 |
| Tecta | 0.287 | 0.041 | N.E.D | |
| Tenm1 | 0.718 | 0.111 | 0.398 | 0.186 |
| Tenm2 | 0.656 | 0.01 | 0.526 | 0.19 |
| Tenm3 | 0.686 | 0.048 | 0.51 | 0.076 |
| Tenm4 | 0.28 | 0.059 | N.E.D | |
| Thsd7a | 0.992 | 0.049 | 0.995 | 0.01 |

**Table S1. 34 extracellular proteins with furin cleavage sites.** A bioinformatics approach identified 11,302 mouse genome proteins with at least one furin cleavage site RX(K/R)R. Using DAVID, these proteins were categorized by extracellular matrix localization, sequence size, and DRG expression patterns via GenePaint [21], revealing 33 proteins with DRG staining signals. Following qPCR quantification of mRNA levels using HRPT as a housekeeping gene (N.E.D., no expression detected), six proteins (highlighted in yellow) were chosen as promising candidates for their specificity to DRG.

Table S2

| Accession | Description | Size (aa) | DRG expression |
|---|---|---|---|
| 8AQT_A | Chain A, Processed angiotensin-converting enzyme 2,Ig gamma-2A chain C region, membrane-bound form | 886 | No |
| 8AQW_A | Chain A, Processed angiotensin-converting enzyme 2,Ig gamma-2A chain C region, A allele | 884 | No |
| 7XMS_S | Chain S, Signle chain viable fragment of antibody | 292 | Not available |
| 6NJL_K | Chain K, 15F1 Fab heavy chain | 262 | Not available |
| 5ZRY_A | Chain A, Ankyrin repeat and SAM domain-containing protein 1A,Ephrin type-A receptor 6 | 194 | Yes |
| 6TFB_A | Chain A, Frizzled-8 | 138 | Yes |

**Table S2. Extracellular proteins with a consensus PreScission cleavage site.** Using a peptide search of the mouse proteome we found six proteins were detected with at least one PreScission consensus cleavage site LEVLFQ/GP.
